# A spatial transcriptomic atlas of autism-associated genes identifies convergence in the developing human thalamus

**DOI:** 10.1101/2025.11.05.685843

**Authors:** Alexander Aivazidis, Fani Memi, Koen Rademaker, Mahmoud Koko, Kenny Roberts, Andrew Trinh, Robert Petryszak, Vitalii Kleshchevnikov, Liz Tuck, Steven Lisgo, Tong Li, Stanislaw Makarchuk, Martin Prete, Tomasz J. Nowakowski, Hilary C. Martin, Omer Ali Bayraktar

## Abstract

Autism is a highly heritable neurodevelopmental condition that manifests across a wide phenotypic spectrum. Rare and *de novo* loss-of-function mutations strongly predispose to autism and co-occurring developmental and intellectual disabilities in over 10% of autistic individuals. Understanding whether these variants converge on specific regional brain circuits or widely alter human brain development is crucial to understanding the etiology of profound autism. To date, transcriptomic atlases have mainly implicated the developing cerebral cortex, yet other brain areas have received relatively little attention. Here, we present a single-cell resolution spatial transcriptomic atlas of 250 autism susceptibility genes during human brain development. Profiling over 10 million cells across the midgestation forebrain, we found convergence of these genes across a small number of regional programs. The developing thalamus showed the most prevalent expression of autism susceptibility genes, followed by germinal zones throughout the brain. Within the thalamus, excitatory neurons showed the most enriched expression, which varied across thalamic nuclei harboring distinct circuits. Across the germinal zones, neural progenitors in the medial ganglionic eminences that generate parvalbumin- and somatostanin-positive interneurons showed highest expression. Our findings reveal the prevalent expression of autism associated genes beyond the developing cerebral cortex and implicate the developing human thalamus as a major hub of autism susceptibility.

## Main

A key question in complex genetic disorders is whether the associated genetic variants exert their phenotypes through convergent cellular and molecular pathways or alter disparate biological pathways. Autism is a highly heritable neurodevelopmental condition that manifests across a wide phenotypic spectrum^1–4^. Large-scale sequencing studies have identified rare and *de novo* loss-of-function mutations in at least 10% of autism cases that strongly predispose individuals to profound autism as well as developmental delay and intellectual disability^5–8^. To date, over 250 genes with rare coding variants that predispose to profound autism have been identified^7–9^. Understanding whether these susceptibility genes affect specific regional human brain circuits or widely alter neural development is crucial to understanding the aetiology of profound autism and choosing the right human model systems, such as region-specific human brain organoids, to interrogate their functions. However, mapping the convergence of autism susceptibility genes across the vast regional cellular heterogeneity of the brain across development^10,11^ poses a major challenge.

A promising approach to infer the spatiotemporal convergence of genetic variants associated with a particular phenotype is to systematically investigate the expression of affected genes in bulk^12^ and single cell^13^ or spatial transcriptomics^14^ atlases of relevant tissues. To date, several bulk and single cell transcriptomics studies have reported the convergence of autism susceptibility gene expression in the mid-gestational prefrontal cerebral cortex, specifically in differentiating excitatory neurons in the developing cortical plate^10,12,15^. However, these studies have largely focused on the cerebral cortex, and have not comprehensively examined other brain regions that could be implicated in autism.

To implicate developing human brain regions and circuits in autism, we mapped the expression of 250 autism susceptibility genes across the mid-gestation human forebrain using single cell-resolution spatial transcriptomics. We found that these genes converged into a small number of highly regionalised expression programs. The developing thalamus emerged as the region with the highest enrichment of autism susceptibility genes, followed by the germinal zones throughout the forebrain. Both regions showed more prevalent autism gene expression than the developing cortical plate and expressed functionally distinct gene sets. Within the thalamus, autism genes associated with neuronal and synaptic functions were enriched in excitatory neurons and their expression varied across thalamic nuclei harboring distinct thalamocortical circuits. Across the germinal zones, autism genes associated with transcriptional regulation showed peak expression in neural progenitors in the medial ganglionic eminences, the primary origin of parvalbumin-and somatostatin-expressing interneurons. Our findings reveal the prevalent expression of autism associated genes beyond the developing cerebral cortex and implicate the developing human thalamus as a major hub of convergence for autism susceptibility.

## Results

### Spatial transcriptomic mapping of autism susceptibility gene convergence

To map where autism susceptibility genes are most highly co-expressed during human brain development, we used the Xenium spatial transcriptomic technology that enables probe-based imaging of single cell transcriptomes *in situ*^*16*^. We curated a Xenium probe panel of 310 unique genes (**Fig. 1A, Supp. Table 1, Methods**). This included 250 autism susceptibility genes (FDR < 0.05) from two recent genetic studies^7,8^ based on rare and *de novo* variants in protein-coding regions. In total, 97 are “high-confidence” autism susceptibility genes with stronger evidence from both studies^7,8^ (FDR < 0.001) and 36 are “autism-predominant” genes (i.e. those more commonly mutated in cohorts for autism rather than for developmental disorders^8^, **Methods**). In addition, our panel included 65 cell type markers spanning neural progenitors and differentiating neuronal and glial cells curated from mid-gestational human brain transcriptome atlases^13,17^, based largely on the cerebral cortex (**Fig. 1A, Supp. Table 1**).

**Figure 1.**
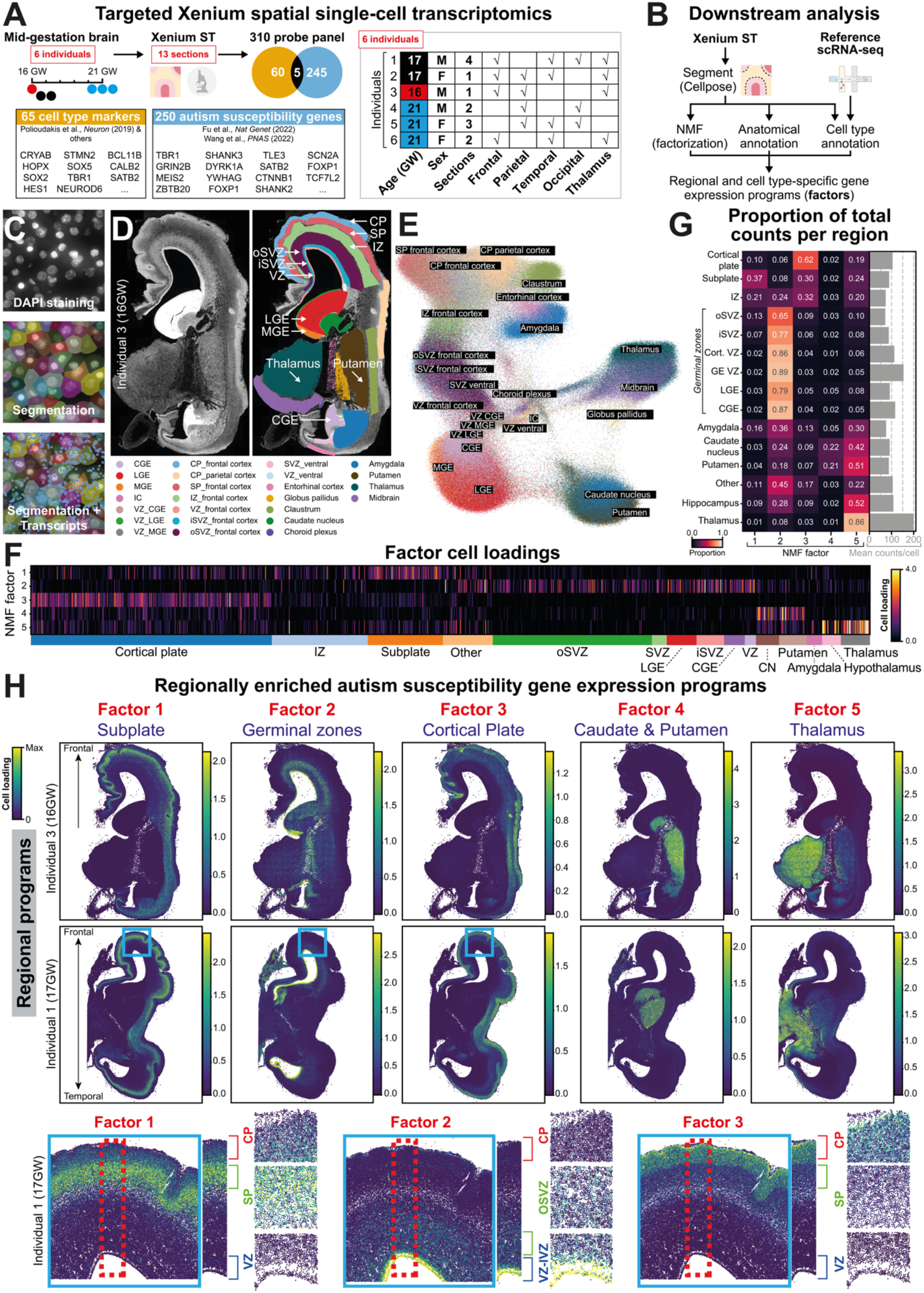
Mapping regional convergence of autism susceptibility genes in the developing human brain using single cell spatial transcriptomics. ***A:*** Xenium spatial transcriptomics with a targeted 310 probe panel of 250 autism susceptibility genes and 65 cell type marker genes (5 overlapping genes, **Supp. Table 1**) was used to assay 13 human mid-gestational forebrain tissue sections from 6 individuals. ***B:*** Analysis strategy to identify regional and cell type-specific autism susceptibility gene expression programs. Following deep learning cell segmentation with Cellpose, single cell autism susceptibility gene expression profiles were factorized using NMF into a small number of convergent gene programs (factors). In parallel, sections were manually anatomically annotated and cell types were classified using Tangram and public reference single-cell RNA-seq data, to interpret regional and cell type specificity of gene programs. **C:** Single cell segmentation of Xenium data. (Top) Nuclei stained with DAPI. (Middle) Cell boundary masks derived from Cellpose nuclei segmentation with 5µm expansion. (Bottom) Subcellular resolution transcripts were counted per segmented cell to derive single-cell level expression data. ***D)*** Anatomical annotation (colour) of one human brain section. Regions were manually labelled based on anatomical landmarks and marker gene expression patterns. ***E)*** Regional segregation of single cell gene expression profiles. UMAP embedding of Xenium transcriptome profiles of single cells derived from panel D. Colours indicate the manual anatomical annotations of individual cells. ***F)*** Regional segregation of autism susceptibility gene programs across single cells. Heatmap shows cell loadings of 5 autism susceptibility gene factors across single cells that cluster by annotated brain regional identity (sidebar). ***G)*** Regional expression pattern of autism susceptibility gene factors. Heatmap shows the proportion of total measured counts explained by each factor per anatomical region. Each factor shows enrichment in at least one region. Barchart on the right shows the mean number of counts per cell per anatomical region. ***H)*** Autism susceptibility gene programs are regionally enriched. (Top two rows) Spatial maps of factor cell loadings across two brains closely track annotated anatomical regions. (Bottom row) Cortical single-cell loadings (close up of area marked by blue box above) highlight specific factor localisation across ventricular zone (VZ), subplate (SP) and cortical plate (CP) in the developing neocortex (marked by red box).

We focused on the mid-gestational human brain (16 to 21 gestational weeks, GW), a period heavily implicated in autism risk^10,12^. We profiled the expression of our 310 genes of interest across 13 large coronal forebrain tissue sections from 6 individuals (3 male and 3 female), containing both cortical and subcortical regions across the rostrocaudal and mediolateral axis (**Fig. 1A, Extended Data Fig. 1, Supp. Table 2**). The Xenium datasets presented here and the associated analysis results below can be interactively navigated via our user-friendly data portal based on WebAtlas^18^ at https://www.stageatlas.org.

To quantify gene expression at single cell resolution, we first segmented nuclei from simultaneously captured DAPI images from Xenium (**Fig. 1B,C**). Comparing three nuclei segmentation approaches, the deep learning model Cellpose^19^ yielded the best results (**Fig. 1C, Extended Data Fig. 2**). For transcript quantification in each cell, we expanded nuclei boundaries by up to 5 μm.

After quality control (**Extended Data Fig. 3, Methods**), our dataset contained 10.86 million single cell gene expression profiles. We then annotated single cells to anatomical brain regions based on histological landmarks and regional marker gene expression (**Fig. 1D, Extended Data Fig. 1**). We observed that single cell transcriptomics segregated by brain regions in sections in the UMAP embedding (**Fig. 1E, Extended Data Fig. 3B**), where cells from germinal zones, cortical plate, subplate and subcortical areas could be distinguished.

To identify whether autism susceptibility gene expression converges on regional programs, we performed Non-Negative Matrix Factorization (NMF) to identify gene programs with correlated expression across cells. We developed a custom NMF implementation that is adapted to technical variation in Xenium experiments, including differences in detection efficiency and background binding across different sections (**Methods, Supp. Comp. Note**). We applied NMF to single cell transcript counts of autism susceptibility genes, excluding cell type markers. This identified 5 factors (i.e. gene programs) with convergent expression of autism susceptibility genes (**Fig. 1F**).

Strikingly, while our NMF approach was unaware of spatial cell locations, we found that gene programs showed highly regionalized expression patterns (i.e. single cell loadings) in the Xenium data (**Fig. 1F-H, Extended Data Fig. 4**,**5**). Autism susceptibility gene factors were enriched in the cortical subplate (factor 1); germinal zones in the cortex and the ganglionic eminences (GEs), including ventricular and subventricular zones (factor 2); the cortical plate (factor 3); the caudate nucleus and putamen (factor 4), and the thalamus (factor 5). These patterns were consistent across multiple individuals (**Extended Data Fig. 6**). Hence, autism susceptibility genes converge on a small number of distinct brain regions in the mid-gestational human forebrain.

### Autism susceptibility gene convergence in the thalamus

Next, we examined the distribution of autism susceptibility gene factors across brain regions. While strongly enriched in one particular brain region, each factor also made minor contributions to autism susceptibility gene expression in other brain areas. For example, factor 5 accounted for 94 counts of susceptibility gene expression (90% of total expression) per thalamic cell on average, yet also accounted for 21 counts of susceptibility gene expression (50% of total expression) per cell in the putamen (**Fig. 1G, Extended Data Fig. 7**).

Given their expression across multiple regions, we quantified the enrichment levels of individual autism susceptibility genes per factor (**Fig. 2A,B**). We defined a *relative gene loading* that quantifies the proportion of mean transcript counts for a gene across anatomically defined brain regions explained by a given factor (**Methods**). We then defined factor-enriched genes as those that show relative gene loading above 33% (**Supp. Comp. Note**). In addition, we classified our panel of 250 autism genes into weakly or robustly expressed, depending on their maximal observed expression across regions (**Methods**). Strikingly, this approach showed that the thalamus factor has the highest proportion of susceptibility genes with 84 robustly expressed factor-enriched genes (**Fig. 2B,C**). This was followed by the germinal zone factor with 49 such genes. The cortical plate, subplate and caudate/putamen factors each included 3-10 robustly expressed factor-enriched genes (**Supp. Table 3**,**4**). The greatest overlap of factor-enriched genes was observed between the thalamus and germinal zones (**Extended Data Fig. 4B**). High-confidence and autism-predominant autism susceptibility genes (**Methods**) occurred at a similar proportion amongst factors (∼40% and 9-14% of genes, respectively) (**Fig, 2B, Supp. Table 3**). Given that NMF is not deterministic, we trained a second NMF model, and this recapitulated the five regionally enriched factors (**Extended Data Fig. 8A-B**) as well as the enrichment of autism susceptibility genes in the thalamus and germinal zones (**Extended Data Fig. 8C**). Furthermore, the distribution of autism genes across factors held up at different relative gene loading thresholds (**Extended Data Fig. 8D**).

**Figure 2.**
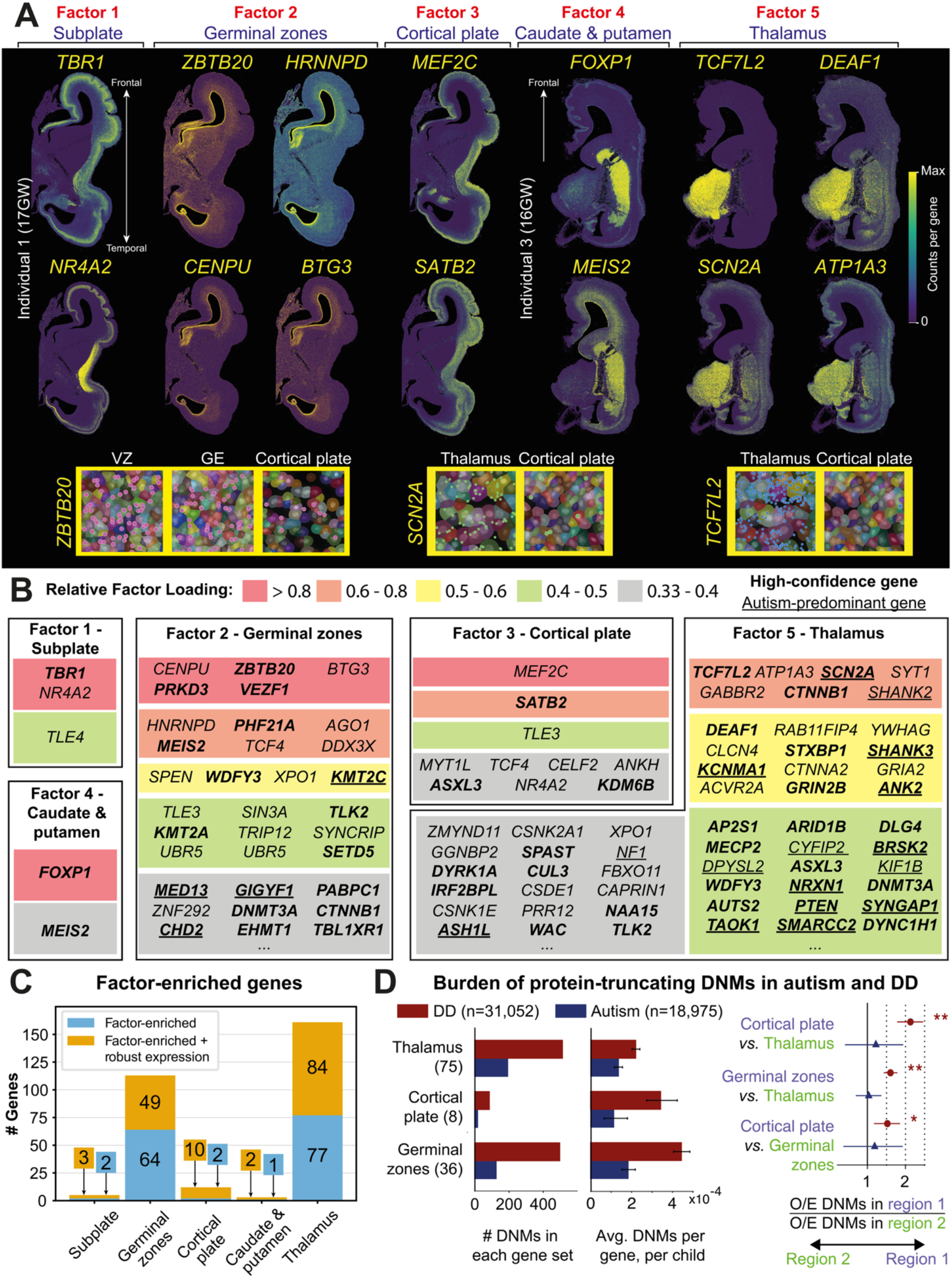
Convergence of autism susceptibility gene expression in the developing human thalamus and germinal zones. ***A)*** Top enriched genes in each factor. Individual genes measured in two human brain sections illustrate the regionalised autism susceptibility gene expression patterns that underlie non-negative matrix factorisation (NMF) results. ***B)*** Table listing factor-enriched autism susceptibility genes. Genes are ordered by their enrichment in each factor from top to bottom (indicated by colour bands) according to relative gene loadings. High-confidence genes are shown in bold and autism-predominant genes are underlined. ***C)*** Number of autism susceptibility genes per factor. A subset of factor-enriched genes (panel B) are genes with robust expression levels across brain regions (orange), remaining factor-enriched genes are shown in blue. The thalamus and germinal zone factors contain the most autism susceptibility genes. ***D)*** Enrichment of protein-truncating *de novo* mutations (DNMs) in robustly-expressed, factor-enriched gene sets. Overlapping genes were assigned to the region with higher factor loading, resulting in thalamus-enriched (n=75), germinal zones-enriched (n=36), and cortical plate-enriched (n=8) gene sets with different numbers than **C**. *(Left)* Total DNM count in the gene sets across DD and autism cohorts. *(Middle)* Average DNM burden per gene, per child in the gene sets across DD and autism cohorts. *(Right)* Pairwise differences in enrichment (observed DNM rate compared to expectation from underlying mutation rates) between brain regions. Arrows indicate the direction of enrichment to a brain region (coloured purple and green). P-values from two-sided Poisson test (^**^ p < 0.05 after Bonferroni correction; ^*^ p < 0.05); error bars show 95% confidence intervals. DD: developmental disorder cohort, O/E: observed over expected ratio. See **Supplementary Note** for details.

As alternative approaches to confirm the regional expression of autism susceptibility genes, we first examined their raw expression in single cells and confirmed the elevated expression of many genes in the thalamus (**Extended Data Fig. 4A**). Second, to quantitatively assess regional convergence, we ranked brain regions by the average number of autism susceptibility genes detected per cell per individual (**Extended Data Fig. 4C**). We used a generalised linear binomial model to test, for each region, if the number of detected autism susceptibility genes (shown in **Extended Data Fig. 4C**) was significantly above the average. This highlighted the thalamus and ventricular zone of the medial ganglionic eminence as enriched for autism susceptibility genes (FDR < 0.001, **Supp. Table 5**). Third, we performed differential expression analysis to extract an orthogonal set of regionally enriched autism genes (**Methods**). While this approach is particularly susceptible to differences in technical detection efficiency within and across Xenium sections, we found broad overlap between the resulting regionally enriched genes and the corresponding factor gene sets (**Supp. Table 6**). Importantly, the thalamus again stood out as harboring 184 regionally differentially expressed genes, followed by the germinal zones with 35 genes. In addition, we detected no significant difference between autism susceptibility gene enrichment between our male and female individuals (**Supp. Table 5**).

Many well-established autism susceptibility genes identified prior to large scale next generation sequencing studies were highly enriched in the thalamus, including *SCN2A*^*20*^, *SHANK2*^*21*^ and *ANK2*^*22*^ (**Fig. 2A,B**). *SCN2A* encodes a voltage-gated sodium channel that propagates excitatory neurotransmission^23^, whereas *SHANK2* and *ANK2* regulate synaptic development and organisation^24,25^. Moreover, SCN2A and ANK2 are known protein interactors^26^. More recently identified autism susceptibility genes were also enriched in the thalamus: *DEAF1* encoding a transcriptional regulator, *ATP1A3* encoding the Alpha-3 subunit of the Na^+^/K^+^ pump^27^ and *GABBR2* encoding a GABAB receptor subunit^28^, as were genes with broad developmental functions, such as *CTNNB1* encoding the key Wnt signalling transducer β-catenin^29,30^ (**Fig. 2A,B**). While expressed across multiple brain regions, other well-studied genes, such as *SYNGAP1* that regulates synaptic development^31^ and *NRXN1* that regulates neuronal connectivity^32^, also showed expression in the thalamus. Hence, the thalamus harbours the expression of well-known and more recently identified autism genes alike.

Top enriched susceptibility genes in the germinal zones included *ZBTB20, PRKD3* and *VEZF1* (**Fig. 2A,B**). Amongst these, *ZBTB20* has a well-characterised role in regulating the timing and differentiation of neural progenitor cells during neurogenesis^33^ and gliogenesis^34^. Across the germinal zones, we noted elevated susceptibility gene expression in GEs, a major source of cortical interneurons^35,36^ (**Fig. 1G**). Cortical interneurons have been implicated in autism by animal studies and post-mortem histological analysis^37^, yet enrichment of autism susceptibility gene expression in the human GEs has not been reported to date. Notably, we observed highest gene expression in the medial GE (**Extended Data Fig. 4A, 9**), the primary source of parvalbumin- and somatostanin-positive interneurons^38^ that have been found to be vulnerable in autism in postmortem studies^39,40^ and mouse models^37^.

In other brain regions, key autism susceptibility genes included *FOXP1* (caudate and putamen), *MEF2C* and *SATB2* (cortical plate), and *TBR1* and *NR4A2* (subplate) (**Fig. 2A,B**). These genes have well characterised roles in the development of their respective regional neuronal populations^41–44^. For example, *TBR1*, enriched in the subplate and developing deep layers, promotes corticothalamic connectivity through the specification of deep layer neuron identity^43^.

We then asked whether rare variants in sets of factor-enriched genes conferred distinct likelihoods for autism and whether this risk is specific to autism or shared with developmental disorders (DDs). Specifically, we examined the burden of protein-truncating *de novo* mutations (DNMs) in robustly-expressed, factor-enriched genes (excluding subplate and caudate putamen genes due to their small sizes) across several published cohorts of individuals ascertained for autism (N=18,975) or for DDs (N=31,052) ^7,8,45^ (**Fig. 2D;** see **Supplementary Note** for details).In autistic probands, the three gene sets showed similar average DNM burden per gene; we found the highest total DNM count in the thalamus gene set followed by the germinal zones, reflecting the size of these gene sets (**Fig. 2D, left**). The three gene sets did not show significant pairwise differences in enrichment (i.e. differences in observed DNM rates) compared to what is expected from their underlying mutation rates (e.g. cortical plate *versus* thalamus, relative risk = 1.17; 95% CI = 0.67 to 1.92; p = 0.50) (**Fig. 2D, right)**. Hence, loss-of-function DNMs in thalamus and germinal zone-enriched genes exert similar degrees of autism predisposition per gene to those enriched in the cortical plate, which is commonly thought to be a hub of autism risk^12,13^. However, due to its higher number of enriched genes, that thalamus factor posed a cumulatively higher risk at the cohort level than other regional factors (**Fig. 2D, left**). Conversely, in DD probands, we found significantly higher burden in the germinal zone (relative risk = 1.52; 95% CI = 1.32 to 1.76; p = 3.7×10^-11^) and cortical plate gene sets (relative risk = 2.19; 95% CI = 1.72 to 2.76; p = 1.0×10^-9^) than in the thalamus (**Fig. 2D, right**). Hence, while loss-of-function DNMs in genes across these regional programs pose similar risk to autism, DNMs in genes enriched in the thalamus pose relatively less risk to DDs than those in germinal zone and cortical plate enriched genes.

Taken together, our results implicate the developing human thalamus as a major hub of autism susceptibility in the developing human brain.

### Autism susceptibility convergence in thalamic excitatory neurons

Next, we examined whether regional autism susceptibility gene expression programs are associated with distinct developmental and cellular functions. First, we performed GO term enrichment analysis of our panel of 250 autism susceptibility genes with all human genes as background and collected enriched terms with at least 5 genes in our panel for further downstream analysis (**Supp. Table 7**). We then classified susceptibility genes into 6 broad functional categories based on enriched GO terms. This resulted in 121 genes assigned to gene expression regulation, 36 genes to axons, 34 to neuronal communication (including synapse formation and ion channels), 23 to the WNT signalling pathway, 23 to cell cycle regulation and 11 to dendrites (**Supp. Table 8**). These results are consistent with previous studies showing autism gene function enrichment in transcriptional regulation and neuronal/synaptic function^6,46^. Across regional programs, we observed enrichment of gene expression regulators in the germinal zones (p < 0.05, Fisher’s exact test, **Methods**) and cortical plate (p < 0.01), corresponding to 68% and 73% of factor genes linked to this category, respectively (**Fig. 3A**). In contrast, only 39% of thalamus factor genes were gene expression regulators (**Fig. 3A**). The thalamus factor harboured more genes in the neuronal communication (p < 0.001), axons (p < 0.001) and dendrite (p < 0.01) categories (18%, 20% and 6% of genes, respectively) compared to the germinal zones (5%, 7% and 1% of genes, respectively) (**Fig. 3A**). These results assign distinct regional functions to spatially expressed autism susceptibility genes in the mid-gestation human brain.

**Figure 3.**
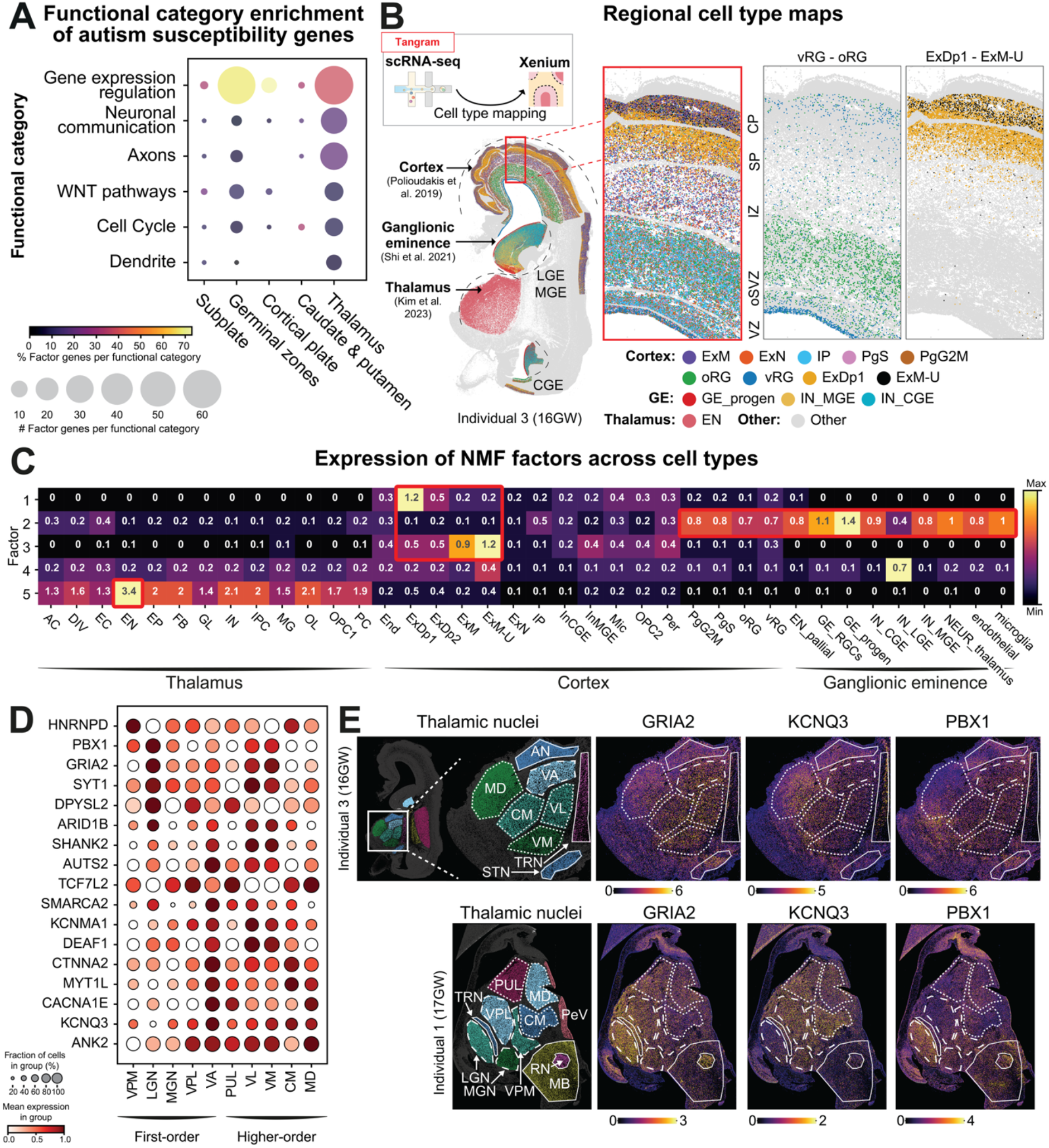
Autism susceptibility gene expression is enriched in developing thalamic excitatory neurons and varies across thalamic nuclei. ***A)*** Functional categorization of factor-enriched genes implicates distinct regional functions. While genes involved in gene regulation make up the largest proportion (dot colour) in all factors, the thalamus factors harbours a relatively lower proportion of these genes and instead included more genes (dot size) involved in neuronal communication (synapse formation, ion channels), axons and dendrites. ***B)*** Regional cell type mapping via integration of matched public single-cell transcriptomics datasets with Xenium spatial transcriptomics. Annotated cortex, ganglionic eminence (GE) and thalamus cells were classified into cell types against public single-cell datasets based on gene expression similarity using Tangram. Top mapped cell types in one 16GW section highlight expected cortical patterning (illustrated in high detail in insets) and diversity in GEs. ***C)*** Heatmap of NMF factor cell loadings across mapped cell types illustrate factor cell loadings converging in specific cell types (marked by red boxes). Subplate (factor 1) and cortical plate (factor 3) map to deep-layer and maturing / upper-layer excitatory neurons, respectively. Germinal zones (factor 2) predominantly map to radial glia and intermediate progenitors across the cortex germinal zones and GE. The thalamus (factor 5) maps to thalamic excitatory neurons. Thalamic cell types^47^: AC, astrocytes; DIV, dividing cells; EN, excitatory glutamatergic neurons; EP, ependymal cells; FB, fibroblasts; GL, glial progenitor cells; IN, inhibitory GABAergic neurons; IPC, intermediate progenitor cells; MG, microglia; OL, oligodendrocytes; OPC1, oligodendrocyte progenitor cells; PC, pericytes. Cortex cell types^13^: Excitatory neuron (EN) types include: ExDp1, deep layer ENs; ExDp2, subplate ENs; ExM, maturing ENs; ExM-U; upper layer ENs; ExN, newborn ENs. End, endothelial cells; IP, intermediate progenitors; InCGE / InMGE, CGE-/ MGE-derived interneurons; Mic, microglia; OPC2, oligodendrocyte precursors; Per, pericytes; PgG2M / PgS, G2M-phase / S-phase cycling progenitors; oRG, outer radial glia; vRG, ventricular radial glia. Ganglionic eminence cell types^35^: EN_pallial, pallial excitatory neurons; GE_RGCs, radial glial cells; GE_prog, intermediate progenitor cells; IN_CGE, CGE interneurons; IN_LGE, LGE interneurons; IN_MGE, MGE interneurons; NEUR_thalamus, thalamic neurons. ***D)*** Expression of genes specific to first-order and higher-order thalamic nuclei in three Xenium sections (**Methods**). ***E)*** Thalamic nucleus-specific expression illustrated in tissue sections from two individuals. Individual thalamic nuclei and other anatomical structures are annotated (left) and spatial gene expression of GRIA2, KCNQ3 and PBX1 are illustrated (right). VPM, Ventral posteromedial nucleus; LGN, lateral geniculate nucleus; MGN, medial geniculate nucleus; VPL, ventral posterolateral nucleus; VA, ventral anterior nucleus; PUL, pulvinar; VL, ventral lateral nucleus; VM, ventral medial nucleus; CM, centromedian nucleus; MD, dorsomedial nucleus, AN, anterior nucleus; PeV, periventricular nucleus; RN, red nucleus; MB, midbrain; TRN, reticular nucleus; STN, subthalamic nucleus.

To determine the cell type specific expression of autism susceptibility genes, we performed cell type classification in our Xenium data in the thalamus, cerebral cortex, and ganglionic eminences using label transfer from published developmental single cell RNA-sequencing datasets of the respective brain regions^13,35,47^ via Tangram^48^ (**Fig. 3B**). We verified expression patterns of known marker genes in corresponding mapped neuroglial cell types in our Xenium data, e.g. *NEUROD6* for cortical neurons, *EOMES* for intermediate progenitors, *HOPX* for outer radial glia and *CRYAB* for truncated radial glia (**Extended Data Fig. 5A**). Maturing cortical upper layer excitatory neurons were the most abundant cell type at more than 1.25 million cells, excitatory neurons represented the largest group in the thalamus with approximately 25,000 mapped cells, and most prevalent in the ganglionic eminences were interneurons with more than 60,000 cells (**Extended Data Fig. 5B**).

We then analysed the average cell loadings of autism susceptibility gene programs in different cell types (**Fig. 3C**). The thalamus (factor 5) showed highest expression in thalamic excitatory neurons (ENs)^47^ (**Fig. 3C**), consistent with neuronal functions of thalamus factor genes described earlier (**Fig. 3A**). Two types of thalamic ENs were previously characterised, and we observed elevated expression in EN2 excitatory neurons, which are enriched for signatures of higher-order thalamic nuclei (see below) (**Extended Data Fig. 5D**). The germinal zones (factor 2) were enriched in progenitors cells, with higher expression in GE radial glia (GE_RGCs) and intermediate progenitors (GE_progen) than cortical progenitors (oRG, vRG, PgS; **Fig. 3C**). Finally, the cortical plate (factor 3) was enriched in excitatory cortical neurons, with elevated expression in maturing (ExM) and maturing upper layer (ExM-U) compared to deep layer neurons (ExDp1, ExDp2; **Fig. 3C**).

Finally, we asked whether autism susceptibility genes associate with distinct neuronal circuits within the thalamus. We compared the gene expression in excitatory neurons across thalamic nuclei, the anatomical subdivisions of the thalamus. Specifically, we compared first-order and higher-order nuclei that are involved in relaying sensory information to primary cortical areas and transthalamic relay of communication between cortical areas, respectively^49,50^. We annotated 10 nuclei across first- and higher-order classes, as well as additional thalamic, midbrain and basal ganglia structures on three 16-17 GW Xenium sections using histological landmarks and regional marker expression (**Methods, Extended Data Fig. 10A,B**). This revealed that most autism susceptibility genes express broadly across multiple thalamic nuclei and without restriction to either first- or higher-order classes (**Fig. 3D, Extended Data Fig. 10C,D**). Yet, we found 17 genes, including 14 thalamus factor-enriched genes, that show regional expression differences. *PBX1, GRIA2, SYT1, ARID1B* and *DPYSL2* showed elevated expression in first-order nuclei and the lateral geniculate nucleus (LGN) in particular, whereas *KCNQ3, DEAF1, SHANK2, ANK2* and *CACNA1E* showed elevated expression across higher-order nuclei (**Fig. 3D,E**). These observations suggest that some autism susceptibility genes associate with distinct regional circuits within the developing thalamus.

## Discussion

Here, we present the first high-resolution spatial transcriptomic atlas of autism susceptibility gene expression in the prenatal human brain to assess the convergence of autism biology across regional brain circuits and cell types. We curated 250 genes in which rare damaging variants strongly predispose to autism and developmental disorders, and measured their expression in over 10 million single cells across different anatomical areas of the midgestation forebrain. An unbiased analysis revealed that these autism genes converge onto five distinct programs. Whereas previous expression profiling studies highlighted cortical excitatory neurons^10,12,15^, our findings unexpectedly highlight the most prevalent autism susceptibility gene expression in thalamic excitatory neurons, followed by neural progenitors across germinal zones.

Our findings challenge the current “cortex-centric” view of autism and implicate the developing thalamus as a major hub of neural circuits likely affected in profound autism. This is consistent with both clinical and neuroimaging evidence. The thalamus and its reciprocal circuits with cerebral cortex are essential for sensory processing and social cognition. Importantly, sensory hyper- and hyposensitivity are amongst core symptoms of autism^51^. The majority of individuals with autism have atypical responses to different types of sensory stimuli^52^, and sensory sensitivities can be amongst the most distressing symptoms for patients, can impair social communication, and predict severity of autism symptoms^53^. Furthermore, neuroimaging studies of individuals with autism have identified complex patterns of thalamic circuit dysconnectivity to cortical areas^54–56^, as early as six months in children with high familial risk for autism^57^.

A major implication of our findings is the need to study autism susceptibility gene function in thalamocortical circuits. The majority of mechanistic work has focused on cortical circuits in animal models and human organoids to date^58,59^. For example, mouse models of *Syngap1* haploinsufficiency show disrupted neuronal maturation and experience-dependent refinement of cortical circuits^60,61^. Similarly, the role of *Scn2a* in synaptic function and plasticity has been mainly studied in the mouse cortex^62^. However, functions of these genes in the developing thalamus have not yet been investigated. In contrast, recent mouse studies of *Shank3* deficiency have found increased excitability of thalamocortical relay neurons^63^ and altered thalamocortical development^64–66^, whereas *Cntnap2* knockout mice show autism-like behaviours linked to reticular thalamic neuron hyperexcitability^67^. Consistently, several autism mouse models show altered somatosensory responses^60,68,69^, and this finding appears to extend to autism individuals and carriers of large copy number variants linked to autism^70^. Functional connectivity interactions between thalamus and the cerebral cortex may involve altered structure or function of thalamic neurons, a finding supported by studies using thalamocortical brain assembloids^71,72^. In addition, we also identified an autism susceptibility gene expression program in the cortical subplate, which guides the development of thalamocortical axons^73^, further pointing at convergent circuit disruption. We propose that both cortical and thalamic circuits are important for the etiology of autism and mechanistic studies of the reciprocal circuits between the two brain regions provides an important area of exploration.

Within the thalamus, we found that autism susceptibility genes are expressed in a varied manner across functionally-distinct thalamic nuclei, implicating excitatory neurons across both first- and higher-order circuits. While our spatial transcriptomic panel did not include extensive markers of fine-grained thalamic neuron subpopulations, we did observe slightly elevated expression in EN2 excitatory neurons that are thought to be more enriched in higher-order nuclei^47^ that relay communication across cortical areas.

Beyond the thalamus, we observed prevalent expression of autism susceptibility gene expression across neural progenitors in the forebrain including both the dorsal VZ and OSVZ as well as ventral GEs. These observations are consistent with mechanistic studies of transcriptional regulators associated with autism in dorso-ventral organoid models^74^. Here we provide the first evidence, to our knowledge, that autism susceptibility genes are enriched in human GEs, a principal source of cortical interneurons. We find elevated expression of these genes in medial GE, the origin of parvalbumin- and somatostatin-positive interneurons, with the former altered in both autism mouse models^37^ and post-mortem autism cases^39,40^, extending the relevance of inhibitory cortical circuits to autism.

The autism susceptibility genes profiled here were compiled from previous genetic studies of autism^7,8^. However, our recent work showed that *de novo* mutations in these genes do not confer sufficient risk on their own to cause autism, although they are sufficiently penetrant to cause DDs (primarily developmental delay and intellectual disability)^45^. While neural progenitors in germinal zones are widely recognized to be altered in DDs^13^, our results suggest that thalamocortical circuits are also broadly relevant to neurodevelopmental disorders. By examining the de novo mutation burden in regionally-enriched autism gene sets across published cohorts, we found here that protein-truncating *de novo* mutations in genes with thalamus enriched expression pose similar risk to autism to germinal zone and cortical plate enriched genes, whereas the former pose less risk to DDs than the latter.

Our study also introduces imaging-based spatial transcriptomics as a promising technology to discover the cellular and tissue contexts of genetic variants. While sequencing-based spatial technologies that profile the whole transcriptome *in situ* have been recently utilised for this purpose^14^, imaging-based technologies such as Xenium^16^ and MERSCOPE^75^ offer 1) true single cell resolution to discriminate cell types (i.e. as opposed to the standard Visium assay) and 2) increased scalability to screen many tissue sections and large tissue areas, enabling better anatomical coverage across complex organs like the human brain. With recent improvements to the number of genes that can be assayed by imaging-based methods^76^, it will be possible to image the whole transcriptome in a routine manner in the future. Finally, the potential integration of spatial transcriptomic maps with brain imaging datasets from individuals with autism^77^ is an exciting future avenue for linking cellular perturbations with circuit-level, cognitive phenotypes.

Our study has several limitations. First, we have mainly focused on autism genes with rare coding variants. While these exert strong effects, common variants genome-wide also play an important role in autism^4,78^, and it has recently been shown that the polygenic basis of autism differs substantially between early- and late-diagnosed individuals^78^. Second, we have mainly focused on midgestation forebrain that is heavily implicated in autism risk. Future work is needed to extend this mapping strategy to other brain regions and developmental periods, in particular early postnatal stages. Last, while we focused on genes that are genetically linked to autism, follow-up mechanistic studies will need to address the function of thalamic circuits in autism.

In conclusion, our work presents the developing human thalamus as a hub of autism susceptibility and a new frontier for autism research.

## Supporting information

Supplementary Computational Note

## Data and code availability

The processed spatial cell atlassing datasets can be explored on our interactive web portal: https://www.stageatlas.org/. The ST datasets will be deposited on the BioImage Archive. The data analysis scripts and notebooks to produce the figures in the manuscript are available here: github.com/BayraktarLab/STAGE. The implementation of NMF for Xenium is available as a Python package: github.com/BayraktarLab/Xenium_NMF.

## Acknowledgements

We thank Aidan Maartens for comments on the manuscript and Kartik Pattabiraman for discussions on thalamic circuits and autism. This work was funded by Wellcome Sanger core funding (220540/Z/20/A) to H.J.M and O.A.B. and a grant from SFARI (736282) to T.J.N. and O.A.B. This work was supported, in part, by the US National Institutes of Health (R01MH125516 to T.J.N.), as well as gifts from the Esther A. & Joseph Klingenstein Fund, Shurl and Kay Curci Foundation, Sontag Foundation, and William K. Bowes Jr. Foundation (to T.J.N.). T.J.N. is a New York Stem Cell Foundation Robertson Neuroscience Investigator. The developing human brain material was provided by the Joint MRC / Wellcome (MR/R006237/1, MR/X008304/1 and 226202/Z/22/Z) Human Developmental Biology Resource (www.hdbr.org).

## Author contributions

*Conception:* A.A., F.M., K.Ra., T.N. and O.A.B. conceived the project.

*Data Generation:* A.A. performed Xenium probe selection, aided by T.N.. F.M. designed Xenium study, selected tissue samples and performed histology, assisted by E.T.. K.Ro and A.T performed the Xenium experiments. S.L. provided tissue samples through the Human Developmental Biology Resource.

*Analysis/Interpretation:* A.A. developed the NMF model. A.A. and K.Ra. performed NMF and downstream analysis. R.P., T.L., S.M. and M.P. developed the STAGE-atlas data portal. T.L. contributed to Xenium segmentation analysis. V.K. contributed to NMF model development. F.M. performed anatomical annotation. M.K. performed enrichment analyses for protein-truncating DNMs across DD and autism cohorts.

*Supervision:* H.M. supervised DNM enrichment analysis. O.A.B. supervised Xenium data generation, analysis and interpretation.

*Manuscript preparation:* A.A., F.M., K.Ra. and O.A.B. wrote the manuscript with feedback from all authors.

## Extended Data Figures and Supplementary Tables

### Extended Data Figures

Extended Data figures 1-10 are provided in an accompanying file.

### Supplementary Tables

Supplementary tables are provided in an accompanying file.

**Table S1 Curated Xenium 310-plex probe panel**

**Table S2 Tissue section and cohort summary**

**Table S3 Summary of enriched genes per factor genes**

**Table S4 Factor-enriched genes and relative gene loading values**

**Table S5 Generalised linear binomial model test of autism susceptibility genes per brain region**

**Table S6 Differential gene expression analysis results of regional marker genes**

**Table S7 Gene Ontology enrichment results for autism susceptibility genes**

**Table S8 Functional categorisation of autism susceptibility genes**

**Table S9 Human mid-gestational brain regional annotation scheme**

**Table S10 Genes from reference single-cell RNA sequencing for Tangram cell type mapping**

## Supplementary information

### Supplementary Note

A Supplementary Note on enrichment of de novo mutations in regionally distinct autism predisposition genes is provided in an accompanying file.

### Supp. Comp. Note

A Supplementary Computational Note on non-negative matrix factorization model adapted for Xenium data is provided in an accompanying file.

## Methods

### Human Tissue

Formalin-fixed paraffin-embedded (FFPE) blocks of second trimester human foetal brain were obtained from the Human Developmental Biology Resource, Newcastle, UK (REC 18/NE/0290).

### Gene panel selection

For our Xenium gene panel, we included all 185 autism susceptibility genes with FDR < 0.05 from their Supplementary Table 11 reported by Fu et al.^8^, which includes rare and *de novo* variants. Our second reference study, Wang et al.^7^, used three models to test for autism associations in *de novo* variants. We included all genes with an FDR < 0.05 in at least two of the three models in their Supplementary Data File 5. We then also included cell type markers from Figure 1F and 2C-D of the Polioudakis et al. single-cell RNA-seq study^13^.

### Definition of autism-predominant and high-confidence autism susceptibility genes

We obtained 36 autism-predominant genes from Supplementary Table 15 in the Fu et al.^8^ study by applying their definition of a posterior probability cutoff >0.99 (“Posterior_probability_ASD” column). We obtained 97 high-confidence autism susceptibility genes as: 72 genes associated with ASD at FDR < 0.001 in their Supplementary Table 11 (“FDR_TADATA_ASD” column) in the Fu et al. study^8^, and genes with FDR < 0.001 in at least one of three statistical models: CH model (“CH_dnLGD_q” or “CH_dnMIS_q” column), denovolyzeR (“DR_dnLGD_q” or “DR_dnMIS_q” column) or DeNovoWEST (“DNW_min_q” column), in Supplementary Data File 5 of the Wang et al. study^7^. Xenium panel genes and their status as autism-predominant and/or high-confidence autism susceptibility genes are reported in **Supp. Table 1.**

### Xenium data generation

FFPE tissue samples were sectioned onto Xenium slides (10x Genomics 3000941) at a thickness of 5 µm and stored at RT in desiccant for up to 4 weeks. These were then processed using the Xenium Sample Preparation Kit (10x Genomics 1000460) as per the manufacturer’s protocols (CG000580 for pretreatment including deparaffinization and decrosslinking, followed by CG000582 for probe hybridisation and rolling circle product generation). Following autofluorescence quenching and nuclei counterstaining as per the manufacturer’s instructions (CG000582), slides were transferred onto a Xenium Analyzer instrument alongside Decoding Reagents (10x Genomics 1000461) and Decoding Consumables (1000487) all prepared according to the manufacturer’s protocol (CG000584). Following the automated assay, in which cycles of hybridisation and imaging are performed in order to assign transcript identities to single RNA spots using a fluorescent barcode readout, data were exported from the instrument in the vendor format of ome.tif images (DAPI-stained nuclei),.h5-format cell-gene count matrices, and.csv-format cell and transcript data (https://www.10xgenomics.com/support/software/xenium-onboard-analysis/latest/analysis/xoa-output-understanding-outputs).

### Brain regional annotation

Brain sections were DAPI- or hematoxylin and eosin (H&E)/Nissl-stained and compared to the Bayer and Altman second trimester human brain atlas^81^ and BrainSpan Reference Atlases for the developing human brain (https://www.brainspan.org/static/atlas). Spatial gene expression patterns from Xenium experiments were inspected and compared against prior regional marker genes to derive anatomical annotations. Absent regional marker genes in the Xenium panel were instead inspected on neighbouring tissue sections that were profiled on 10x Genomics Visium (unpublished data). In short, *CRYAB* and *PDGFD* were used to differentiate cortical VZ from SVZ^82^; *HOPX, GLI3, SOX2* and *EOMES* to mark the extent of oSVZ^82^; *NR4A2*/*NURR1* for subplate^83,84^ and claustrum^85,86^, *NKX2*.*1* and *LHX6* for MGE, and *SP8* for LGE and CGE^87^. The thalamus was identified by the expression of *TCF7L2* ^47,88,89^.

In total, 10,858,858 cells were regionally annotated to 62 regions across 12 systems (e.g. cortex, ganglionic eminence, thalamus) and orientations (e.g. frontal, temporal, occipital, parietal) (**Supp. Table 9**). In addition to this, we annotated 10 thalamic nuclei in three relevant Xenium slides: individual 1 (Sob23-BRA-0-FFPE-5), individual 2 (Sob20-BRA-0-FFPE-4-S25) and individual 3 (Hob5-BRA-5-FFPE-1-S142) (**Supp. Table 2, Supp. Table 9**). Matching H&E stains were annotated for thalamic nuclei using the Bayer and Altman second trimester human brain atlas, and annotations were transferred to segmented Xenium cells. Thalamic nuclei were categorised into first-order and higher-order nuclei based on existing literature^90–92^, and we additionally annotated other thalamic structures and non-thalamic structures (**Extended Data Fig. 10**). Thalamic nuclei that are composed of a mixture of first-order and higher-order subdivisions were categorised into other thalamic regions, and included the thalamic reticular nucleus (TRN)^90,93^. We restricted analysis to 19545, 6717 and 20331 cells classified with Tangram as thalamic excitatory neuron subtypes EN1 or EN2, as defined by Kim et al.^47^ for individuals 1-3, respectively (**Extended Data Fig. 10E**). To assess elevated gene expression in particular nuclei (**Fig. 3D**), raw count data were normalised to 1000 counts per cell (function *sc*.*pp*.*normalize_total*) and ln(x+1) transformed (function *sc*.*pp*.*log1p*), followed by ranking genes per thalamic nuclei (function *sc*.*tl*.*rank_genes_groups*, Wilcoxon rank-sum test), and finally spatial expression of identified genes was visually validated using the WebAtlas portal: https://www.stageatlas.org/.

### Xenium cell segmentation

We benchmarked alternative segmentation methods Cellpose^19,94^ and Proseg^95^ against default on-instrument nucleus segmentation and expansion (Xenium Onboard Analyser or “XOA” hereafter) with custom scripts. Default nuclei segmentation masks (XOA versions 1.4, 1.6, or 1.7, see **Supp. Table 2**) had been expanded by 15µm (pixel size of 0.2125µm) or until encountering other boundaries (output name “xenium_15_micron”). We manually expanded by 5µm from “xenium_15_micron” nuclei masks and refer to this segmentation as “xenium_5_micron”. This expansion has become the default setting from XOA version 2.0 onwards. Cellpose model ‘cyto3’ (version 3.0.3) was used to segment nuclei with 25 pixel cell diameter and was followed by 23 pixel expansion (“cellpose_5_micron”). Transcripts were reassigned to cells and X/Y cell centroid coordinates were extracted with Python package scikit-image (version 0.23.2) function *skimage*.*measure*.*regionprops*. Proseg segmentation (version 1.0.3) was performed on Xenium transcripts with *--xenium* flag and default parameters (‘proseg’).

Cell-gene count matrices were generated in AnnData (version 0.10.6) format with Scanpy^96^ (version 1.10.3) for all segmentation method outputs. We quantified the following cell segmentation metrics using Github repository ‘Xenium_benchmarking’^97^ (https://github.com/Moldia/Xenium_benchmarking, commit 1908b7f at 30 June 2024): proportion of assigned reads, number of identified cells, reads per cell (median, 5th and 95th percentiles), genes per cell (median, 5th and 95th percentiles), cellular density (number of cells / section area in µm^2^).

We did not strictly optimise a segmentation strategy to maximise proportion of reads assigned to cells, since xenium_15_micron inherently assigned more reads compared to other strategies with only 5µm expansion. Instead, we opted for Cellpose because of positive quantitative metrics (**Extended Data Fig. 2**) with fewer total cells, yet higher median transcript and gene coverage. Furthermore, we conducted manual visual inspections of dense tissue regions such as germinal zones and upper cortical layers to further confirm segmentation performance.

### Xenium data processing and quality control

Cell-gene count matrices were processed with Scanpy (v1.10.3) following transcript reassignment to Cellpose cell segmentation masks. We evaluated cell metrics including transcripts and genes per cell, and cell area (**Extended Data Fig. 3A**), and we filtered out cells with high transcript numbers (>1000) as these could represent doublets.

Additionally, raw count data were normalised per cell (function *sc*.*pp*.*normalize_total*) and ln(x+1) transformed (function *sc*.*pp*.*log1p*), and together with the nearest neighbor graph (function *sc*.*pp*.*neighbors*, n_neighbors=15) passed to the function *sc*.*tl*.*umap* (min_dist=2.0, spread=2.0) to calculate a Uniform Manifold Approximation and Projection (UMAP) embedding^98^. Brain regional information and tissue section were projected onto the UMAP embedding (**Extended Data Fig. 3B-C**).

### Non-negative matrix factorization

We used a customized implementation of non-negative matrix factorization (NMF) in the probabilistic programming language Pyro^99^ that included parameters to model technical factors of variation that arise in Xenium experiments.

Non-negative matrix factorization decomposes the matrix of biological expression value 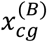 of size cells, *c*, and genes, *g*, into two smaller matrices of dimensions cells x factors and factors x genes^100^:

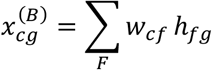

We take into account technical factors of variation that change the expectation values for measured quantities of counts by transforming “biological” expectation values to account for technical variables:

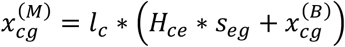

Here, *H*_*ce*_ denotes a one-hot categorical assignment of cells to experimental batches - which correspond to individual sections in Xenium data - and *l*_*c*_ describes differences in detection efficiency of genes across cells, and *s*_*eg*_ models ambient RNA (e.g. “background binding”) for each gene in each batch. These “measurement” expectation values are then used to parameterize a Negative Binomial observation model of the observed raw count values *X*_*cg*_:

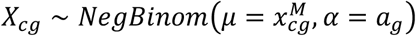

Model training was performed on an NVIDIA A100-SXM4-80GB GPU using the Pyro stochastic variational inference framework^99^ for 40,000 epochs with a learning rate of 0.001, a batch size of 2,171,772 cells and used 10 initial factors. After training we plotted the 99.9% quantile of gene and cell loadings of factors. This revealed a cluster of 5 ‘active’ factors with high loadings and a cluster of ‘inactive’ factors with low loadings, which we removed for downstream analysis (**Extended Data Fig. 8E-F**). To assess the stability of our NMF results, we also performed a second NMF training run with identical parameters yet different initialisation and random number seeds, which again resulted in 5 active factors. Region-specific expression of 3 factors (1: cortical plate; 2: germinal zones; 5: subplate) recapitulated similar patterns to the first training run (**Extended Data Fig. 8B**). However, factor 4 merged the previous factor with lowest counts (i.e. caudate nucleus and putamen in the basal ganglia) into the previous factor with highest counts (expressed in thalamus). Finally, factor 5 was observed across basal ganglia and other structures, yet not specifically in the thalamus. We kept the first NMF training run for downstream results because it provided a more interpretable region-specific expression of factors (**Extended Data Fig. 8A**), and a better mean absolute error fit of predicted expression values to observed Xenium counts.

More information on prior distributions of all free parameters, the inference procedure, initialisation of factors, and the definition of relative gene loadings is included in the **Supp. Comp. Note**).

### Robust expression threshold

Across 6,912,281 cells in the cortical CP, SP, IZ, iSVZ/oSVZ, VZ; ganglionic eminences MGE, LGE, CGE and thalamus (excluding lateral geniculate nucleus) (**Supp. Table 9**), the mean expression of genes was calculated in each region. The maximum mean expression across regions was calculated for each gene (5-95% quantile 0.023-3.375, 95%, min. 0.014, max. 12.196 counts), and the minimum threshold for robust expression was set to the median (0.55) of maximum mean expression of all genes (**Supp. Table 4**).

### Xenium cell type annotation

We leveraged three publicly available scRNA-seq reference datasets (**Supp. Table 10**) for cell type label transfer onto Xenium cells with Tangram^48^ (version 1.0.4): human developmental neocortex at 17-18 GW^13^, ganglionic eminence at 9-18 GW^35^ and thalamus at 16-25 GW^47^. Neocortex and thalamus datasets already included coarse and granular cell type annotations. We re-analysed ganglionic eminence data in Scanpy to mimic author instructions for the R language package Seurat. Briefly, counts were normalised (*sc*.*pp*.*normalize_total* function, target_sum=1000) and log1p-transformed (*sc*.*pp*.*log1p* function). An scVI ^101^ (version 1.0.4) variational autoencoder was trained on 2000 most highly variable genes (*sc*.*pp*.*highly_variable_genes* function, flavor=‘seurat’, min_mean=0.1, max_mean=8) with 30 latent variables and 2 hidden layers with 1024 nodes each, under a negative-binomial gene likelihood distribution to integrate expression data from multiple samples. This latent representation was used to compute the neighbourhood graph (*sc*.*pp*.*neighbors* function with 10 neighbours), Leiden clustering^102^ with resolution=0.3, UMAP (min_dist=0.3, n_components=2), and we scored cluster marker genes (*sc*.*tl*.*rank_genes_groups* function, Wilcoxon rank-sum test). This recapitulated 9 clusters: MGE (LHX6^+^), GE progenitors (MKI67^+^, HMGN2^+^), CGE (NR2F1^+^), LGE (MEIS2^+^), pallial cells (NEUROD2^+^), thalamic neurons (TCF7L2^+^, LHX9^+^), GE RGCs (PTN^+^, TTYH1^+^, FABP7^+^), microglia (CD68^+^), endothelial cells (IFGBP7^+^).

We mapped both coarse and granular cell types to expected annotated anatomical regions, e.g. the developmental neocortex dataset only to annotated cortical regions. Xenium count data were log1p-normalised (*sc*.*pp*.*normalize_total* and *sc*.*pp*.*log1p* functions with default parameters) to match scRNA-seq data as a requirement for Tangram. Intersecting and non-zero expressed genes from both modalities were used as training sets: with 299, 66 and 308 genes for cortex, ganglionic eminence and thalamus respectively (**Supp. Table 10**). We trained Tangram for each tissue section-segmentation combination with default learning rate (0.1) for 1000 epochs, in ‘clusters’ mode with a uniform density prior, and transferred cell type labels to Xenium cells from output probabilistic mapping.

### GO enrichment

We first performed GO term enrichment analysis of our panel of 250 ASD susceptibility genes using the GSEApy Python package ^103^ (version 1.1.8) with all human genes as background. We collected enriched terms (Bonferroni p-value below 0.05) with at least 5 genes in our panel for further downstream analysis (**Supp. Table 7**). We then classified these terms into 6 broad functional categories, including gene expression regulation, axons, neuronal communication (including synapse formation and ion channels), WNT signalling pathway, cell cycle and dendrites (**Supp. Table 8**) and then assigned genes to one or multiple of these categories, based on their GO term annotations. 76 autism susceptibility genes were not assigned to a category, because they had less specific, less frequent or no enriched GO term assigned to them.

### Enrichment of functional categories in factor genes

We used Fisher’s exact test to test the null hypothesis that the proportion of genes annotated with a functional category is the same in each set of factor genes of interest as in the remaining set of genes. The total background set of genes was all ASD susceptibility genes in our panel. The total background set of genes for each functional category was given by the total number of genes annotated with this functional category.

### Differential Gene Expression Analysis

We used the rank-sum test on raw single-cell counts between each region and the rest with the Scanpy (v1.10.3) function *sc*.*tl*.*rank_genes_groups* to extract regional marker genes. We kept genes with an adjusted p-value above 0.05. We subset cells to 25000 per region, equally spread across sections, both to reduce the memory requirements of this analysis and to prevent regions from having more marker genes only because they are overrepresented in our dataset (e.g. there are many more cells from the cortex than the putamen).

### Linear Binomial Model

Following a risk gene enrichment enrichment approach from a previous single-cell RNAseq atlas^13^, we used a generalized linear binomial model to model the number of detected autism susceptibility genes in each cell, with brain region identity and sex as covariates. We used the Python statsmodels package (version 0.14.4) to compute the effect sizes, p-values and FDR for all covariates, which are included in **Supp. Table 5.**

