## Extended Figures and Supplementary Note for "A spatial transcriptomic atlas of autism-associated genes identifies convergence in the developing human thalamus"

### Extended Data Figures

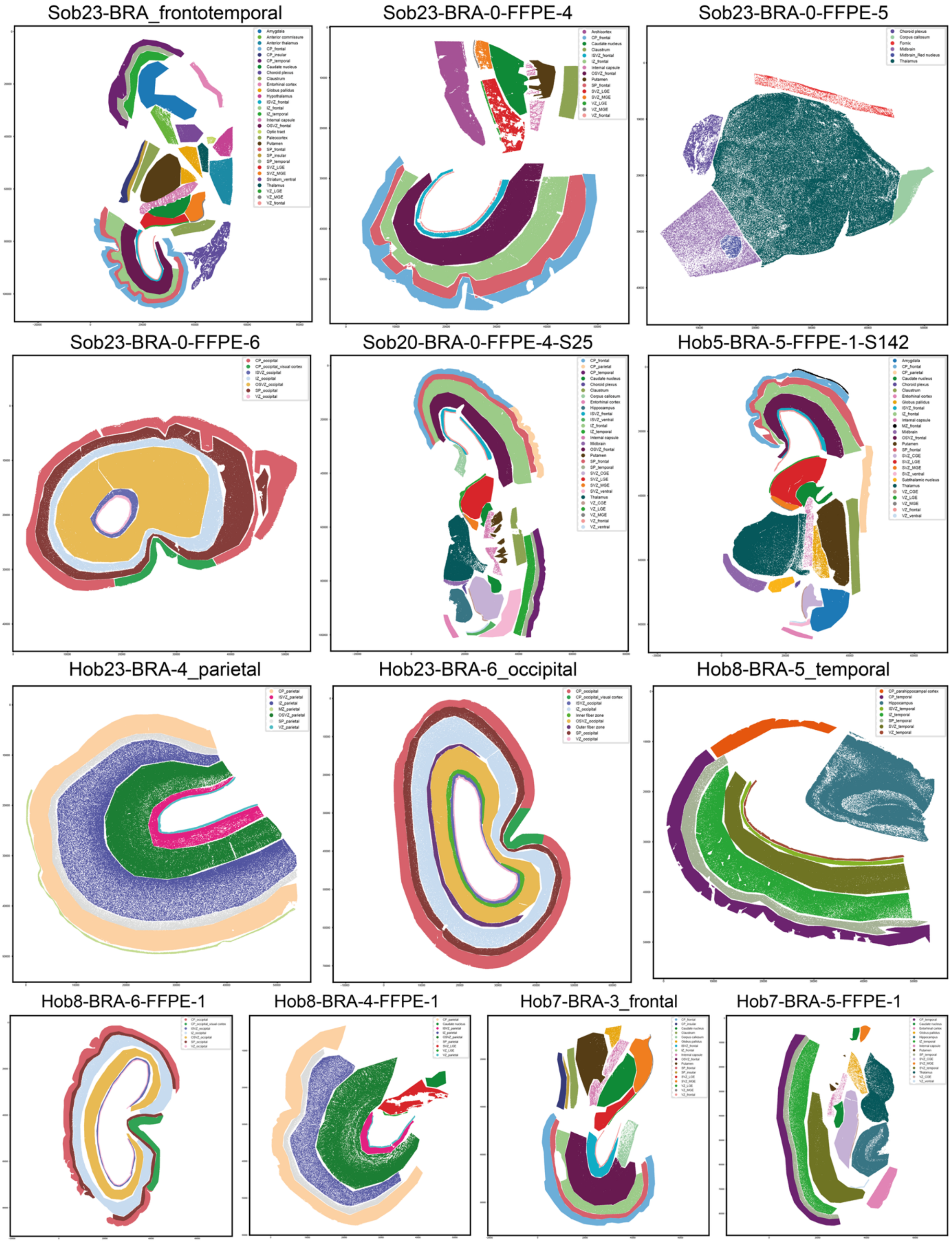


***Extended Data Fig. 1: Overview of mid-gestational brain tissue sections and their anatomical annotation used for Xenium profiling.***

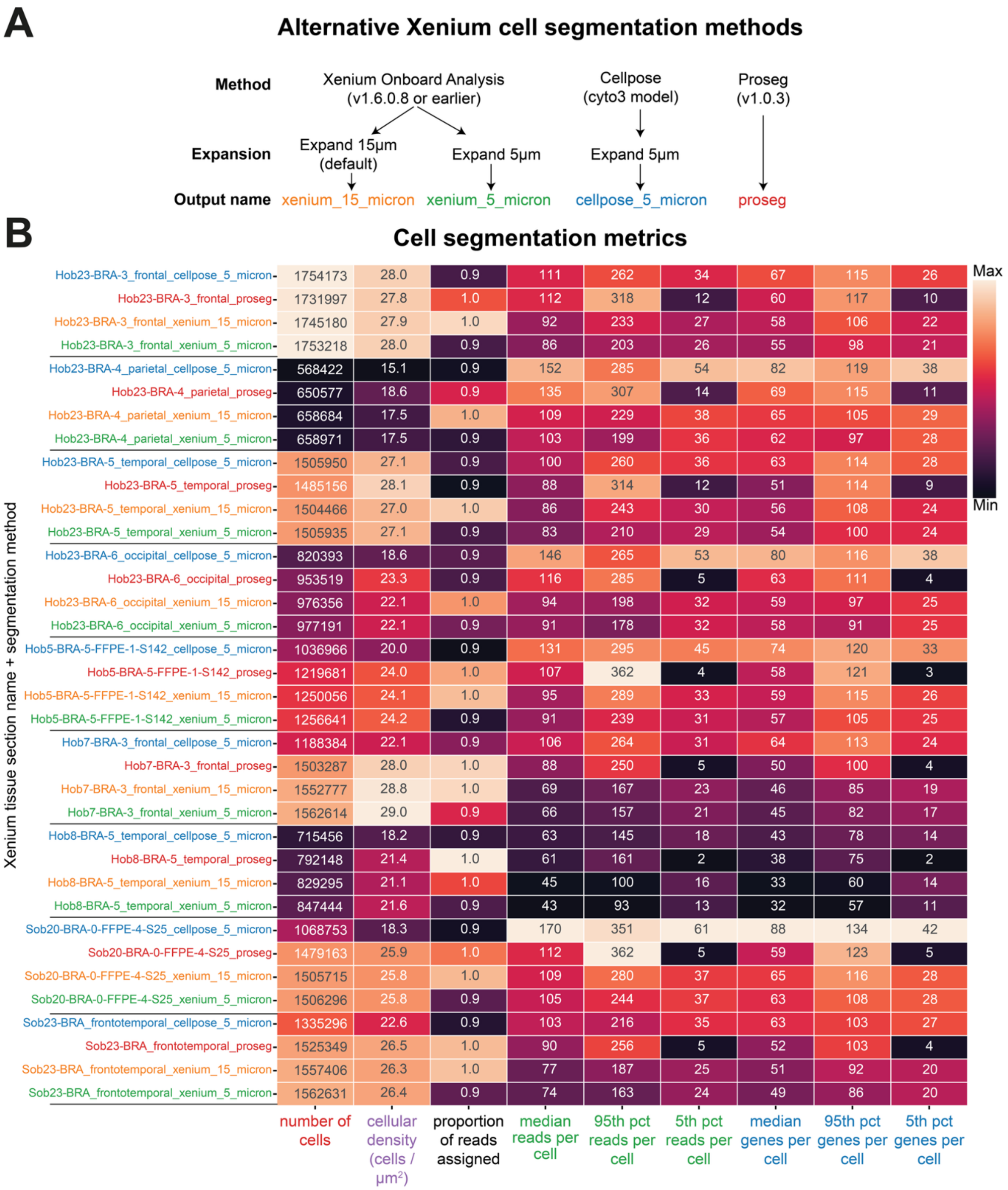


***Extended Data Fig. 2: Comparison of 3 different nuclei segmentation pipelines based on QC metrics***

***A)*** Workflow of alternative cell segmentation methods. Xenium Onboard Analyser, Cellpose and Proseg were run for each tissue slide to segment DAPI-stained nuclei, followed by optional expansion of 15μm or 5μm, to obtain cell masks. Default on-instrument segmentation at time of the experiments was Xenium Onboard Analysis with 15μm expansion.

***B)*** Alternative cell segmentation method performance. Heatmap represents the performance of segmentation methods on tissue slides (rows) with quality metrics (columns) such as number of segmented cells, cellular density, proportion of reads assigned, statistics for reads per cell and statistics for genes per cell. Methods includE) Xenium Onboard Analyser with 15μm expansion (“xenium_15_micron”), Xenium Onboard Analyser with 5μm expansion (“xenium_5_micron”), Proseg (“proseg”), Cellpose cyto3 model with 5μm expansion (“cellpose_5_micron”).

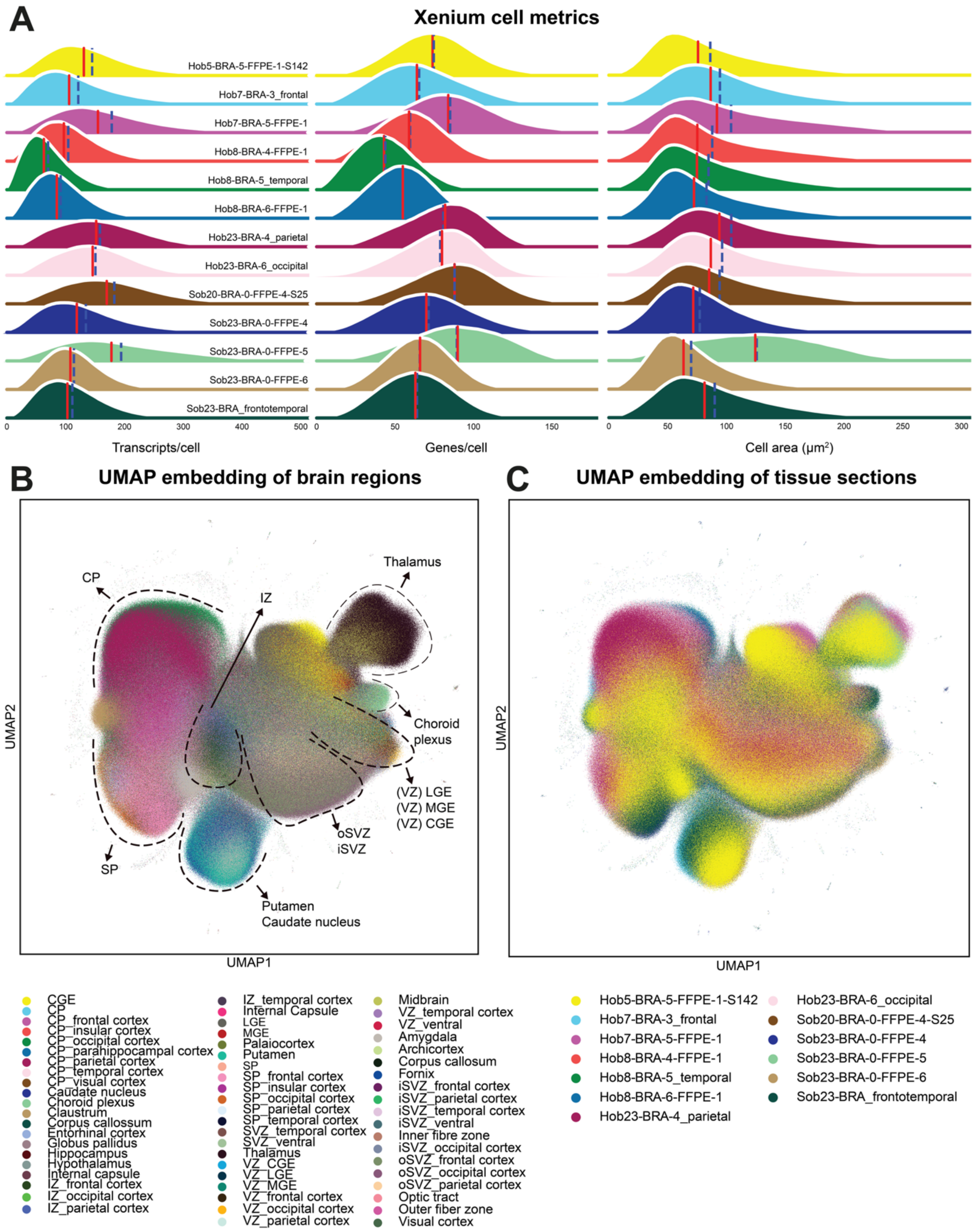


**Extended Data Fig. 3: Overview of Xenium quality control and UMAP embeddings.**

***A)*** Density plots of mean (blue line) and median (red line) Xenium cell metrics, including transcripts per cell (left), genes per cell (middle) and cell area in μm$^2$ from Cellpose segmentation (right) for each named tissue section (rows).

***B)*** UMAP projection of single-cell spatial gene expression per annotated brain region. Dotted lines indicate brain regions into which single-cell transcriptomes broadly segregate, including subplate (SP), cortical plate (CP), intermediate zone (IZ), basal ganglia and its components, outer- and inner subventricular zone (oSVZ, iSVZ), thalamus, and the lateral, medial and caudal ganglionic eminences (LGE, MGE, CGE) and their ventricular zones (VZ).

***C)*** UMAP projection of single-cell gene expression per tissue section.

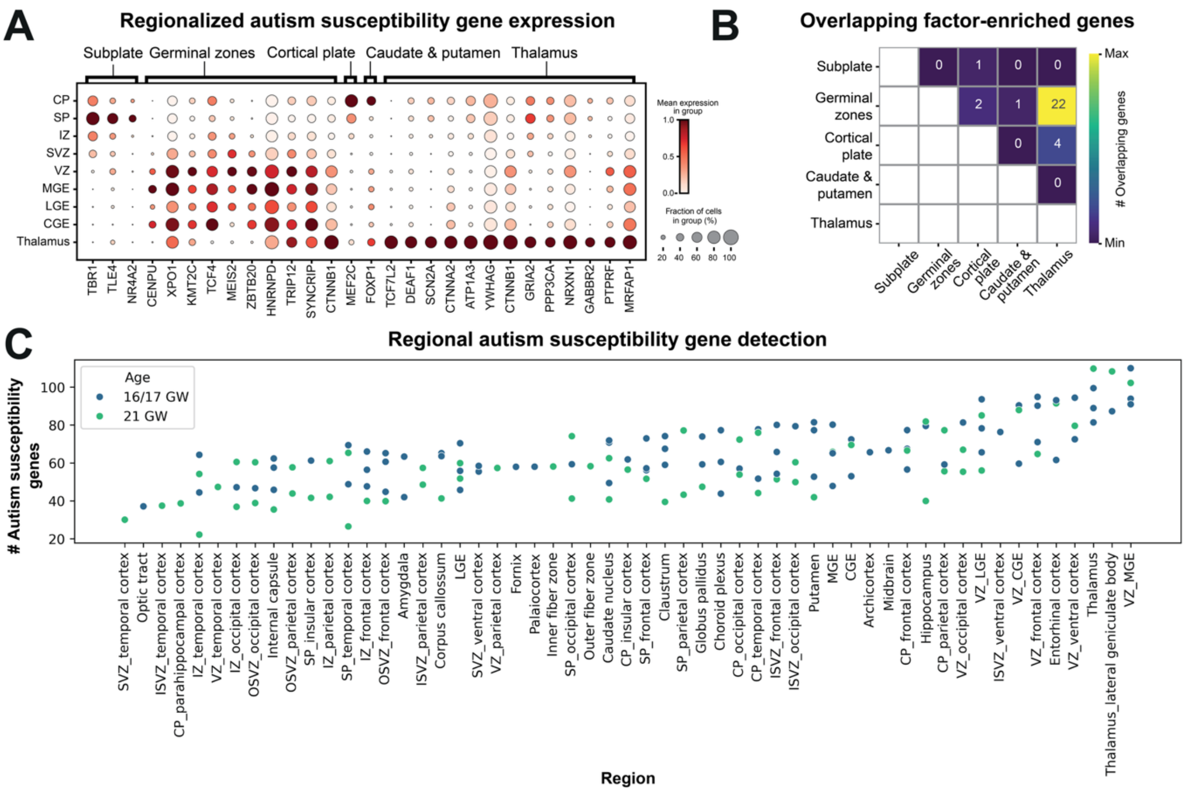


**Extended Data Fig. 4: Regional autism susceptibility gene expression and detection.** ***A)*** Expression of top factor genes across brain regions. Genes such as SCN2A that have been functionally investigated in the cortex, exhibit higher expression in the thalamus. ***B)*** Overlap in factor-enriched genes (includes both weakly or robustly expressed genes). The thalamus and germinal zones factor share most (22) autism susceptibility genes. ***C)*** The average number of autism susceptibility genes detected in cells of each subregion for each individual at age groups 16/17 or 21 GW (gestational weeks), highlights the thalamus and ventricular zone of the medial ganglionic eminence.

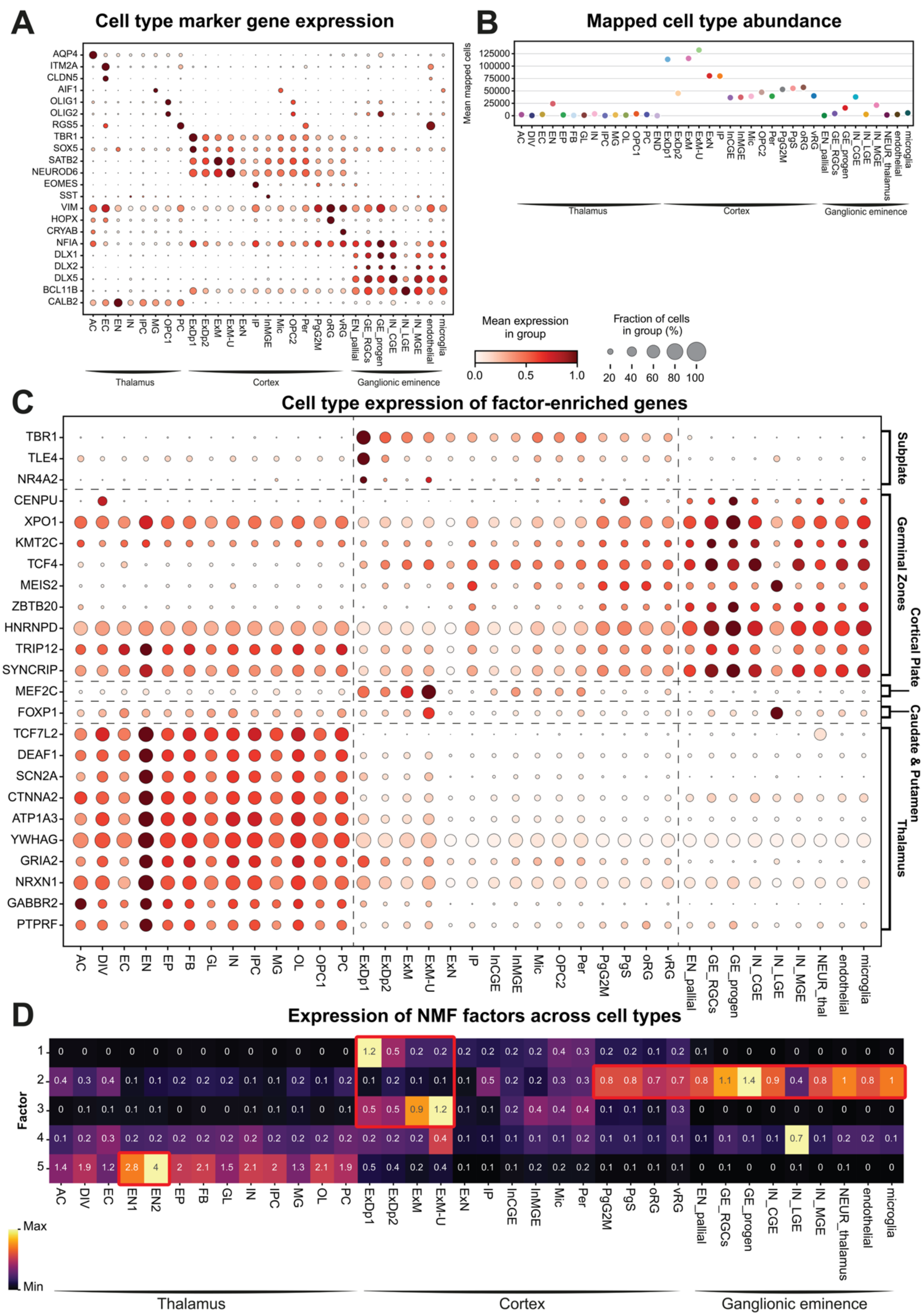


**Extended Data Fig. 5: Brain regional cell type mapping of Xenium cells.**

***A)*** Expression of literature-derived marker genes for mapped thalamus, cortex and ganglionic eminence cell types in Xenium cells. Cells were mapped per annotated brain region to corresponding reference single-cell RNA-sequencing datasets with Tangram.

***B)*** Abundance of Tangram regional mapped cell types averaged across individuals. Cortical cell types were mapped most, followed by ganglionic eminence types and finally excitatory thalamic neuron (EN).

***C)*** Cell type expression of factor-enriched genes. Factor-enriched genes (left, grouped per factor on the right) display specific expression in specific and relevant cell types, e.g. cortical excitatory neurons (ExDp1, ExDp2, ExM and ExM-U) in the subplate and cortical plate factors.

***D)*** Extended heatmap of NMF factor cell loadings illustrate factor cell loadings converging in specific cell types (marked by red boxes). Highlighted is the slightly elevated expression in EN2 compared to EN1 excitatory neurons compared in the thalamus (factor 5). Subplate (factor 1) and cortical plate (factor 3) map to deep-layer and maturing / upper-layer excitatory neurons, respectively. Germinal zones (factor 2) predominantly map to radial glia and intermediate progenitors across the cortex germinal zones and GE.

Thalamic cell types[47]: AC, astrocytes; DIV, dividing cells; EN1/2, excitatory glutamatergic neurons; EP, ependymal cells; FB, fibroblasts; GL, glial progenitor cells; IN, inhibitory GABAergic neurons; IPC, intermediate progenitor cells; MG, microglia; OL, oligodendrocytes; OPC1, oligodendrocyte progenitor cells; PC, pericytes.

Cortex cell types[13]: Excitatory neuron (EN) types include) ExDp1, deep layer ENs; ExDp2, subplate ENs; ExM, maturing ENs; ExM-U; upper layer ENs; ExN, newborn ENs. End, endothelial cells; IP, intermediate progenitors; InCGE / InMGE, CGE- / MGE-derived interneurons; Mic, microglia; OPC2, oligodendrocyte precursors; Per, pericytes; PgG2M / PgS, G2M-phase / S-phase cycling progenitors; oRG, outer radial glia; vRG, ventricular radial glia.

Ganglionic eminence cell types[35]: EN_pallial, pallial excitatory neurons; GE_RGCs, radial glial cells; GE_prog, intermediate progenitor cells; IN_CGE, CGE interneurons; IN_LGE, LGE interneurons; IN_MGE, MGE interneurons; NEUR_thalamus, thalamic neurons.

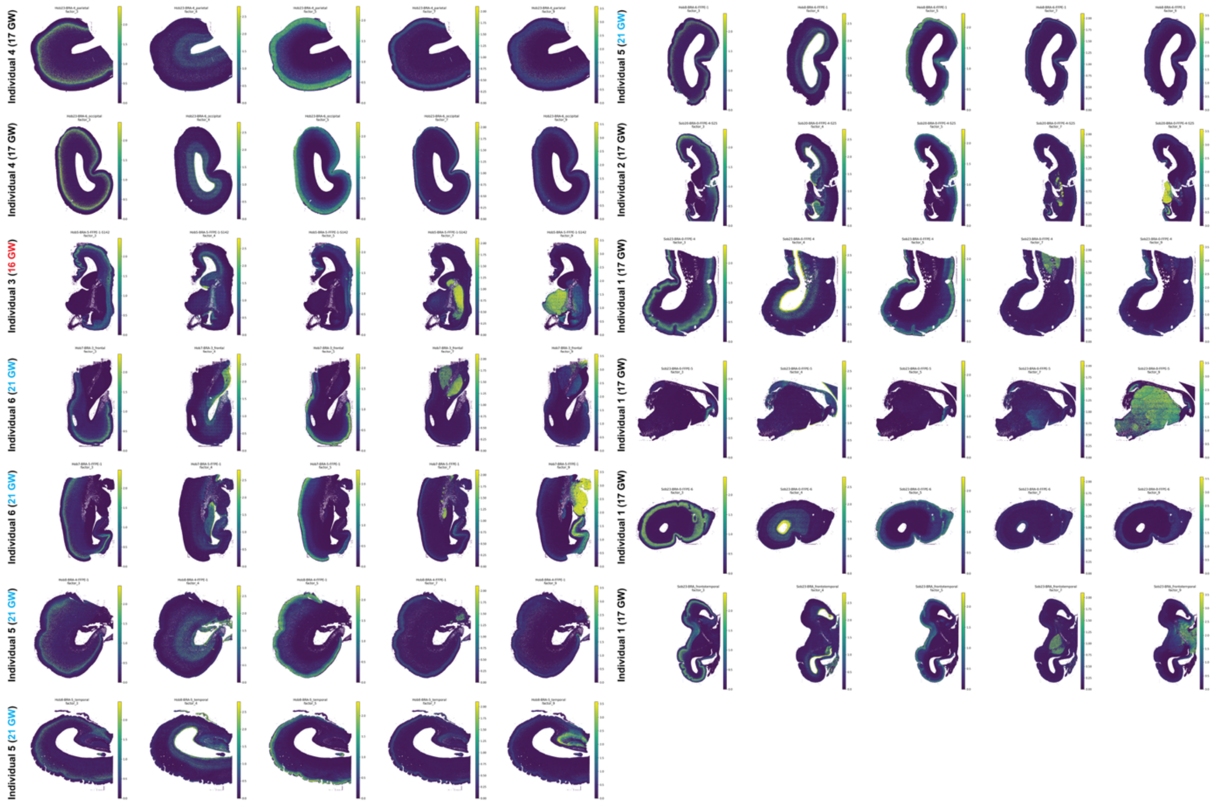


**Extended Data Fig. 6: Non-negative matrix factorization (NMF) single-cell factor cell loadings.**

Cell loadings for NMF factors 1-5 (left to right columns) per mid-gestational brain tissue section (rows). Factors correspond to: cortical subplate ("factor_3"), the germinal zones in the cortex and the ganglionic eminences including ventricular and subventricular zones ("factor_4"), the cortical plate ("factor_5"), the caudate nucleus and putamen ("factor_7") and the thalamus ("factor_9").

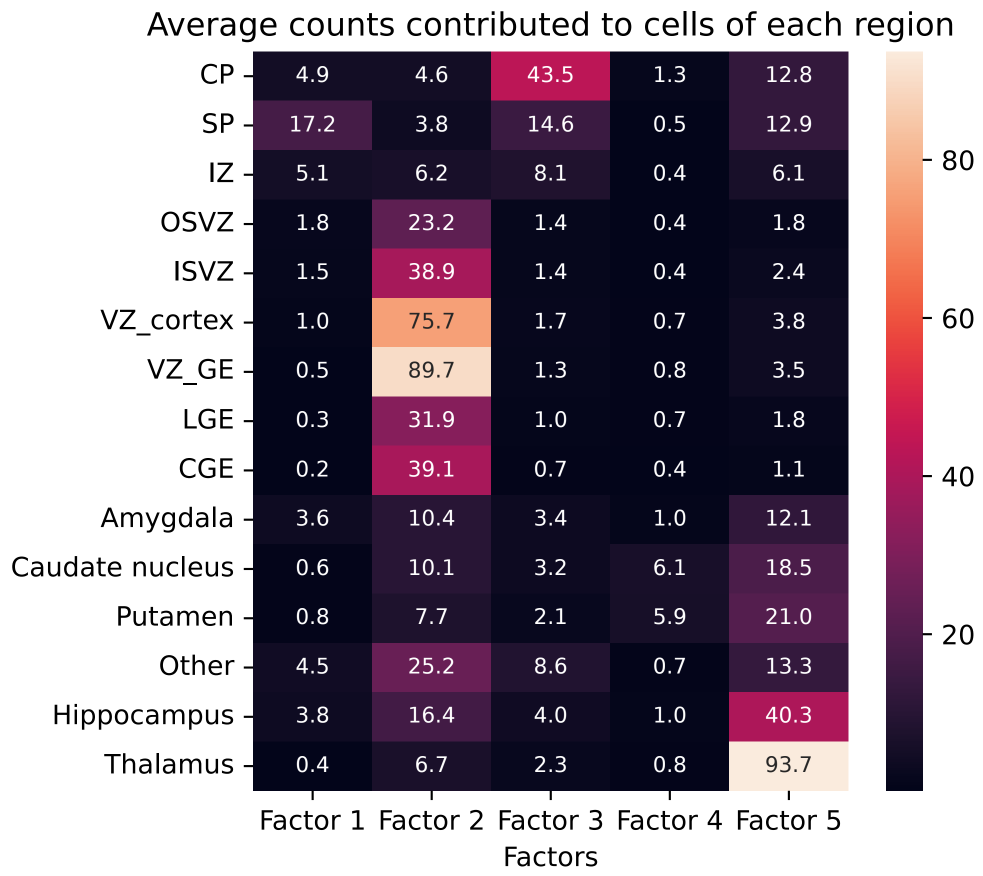


**Extended Data Fig. 7: Average cell counts of NMF factors in each brain anatomical region.**

Average counts contributed to cells in each region by each factor. Factor 2 (germinal zones) and 5 (thalamus) contribute the largest number of counts, while factors 4 (caudate nucleus and putamen), factor 1 (cortical plate) and factor 2 (subplate) only contribute a small number of counts.

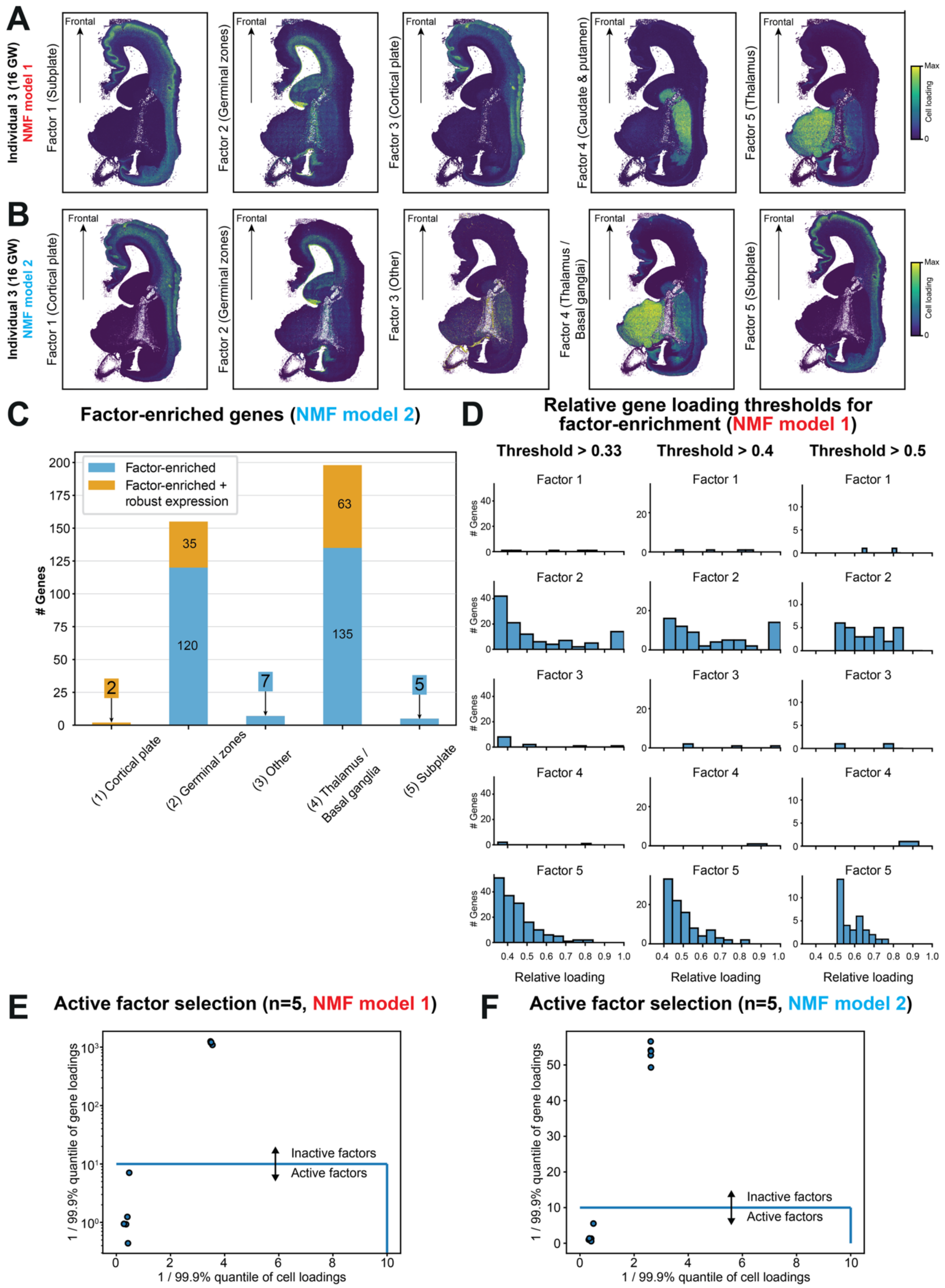


**Extended Data Fig. 8: Comparison of two non-negative matrix factorization NMF training runs.**
***A)*** Spatial maps of factor cell loadings from NMF model 1 highlight localisation to anatomical regions: (1) subplate, (2) germinal zones, (3) cortical plate, (4) caudate nucleus and putamen, and (5) thalamus.

***B)*** Spatial maps of factor cell loadings from NMF model 2 highlight distinct localisation: (1) cortical plate, (2) germinal zones, (4) thalamus and basal ganglia, (5) subplate, (3) other regions including basal ganglia.
***C)*** Active factor selection for NMF model 1 derived by thresholding (indicated by arrows) the 99.9% quantiles of gene and cell loadings highlights 5 factors.
***D)*** Active factor selection for NMF model 2 derived by thresholding (indicated by arrows) the 99.9% quantiles of gene and cell loadings highlights 5 factors.
***E)*** Alternative relative gene loadings thresholds to define factor-enriched genes for NMF model 1. A threshold of 0.33 was previously derived (**Supp. Comp. Note**). Patterns of factor-enriched gene numbers hold up at higher threshold values of 0.4 or 0.5.
***F)*** Number of factor-enriched and robustly expressed genes for NMF model 2 identifies the highest number of genes in the thalamus and germinal zones, similar to NMF model 1 (shown in **Fig. 2C**).

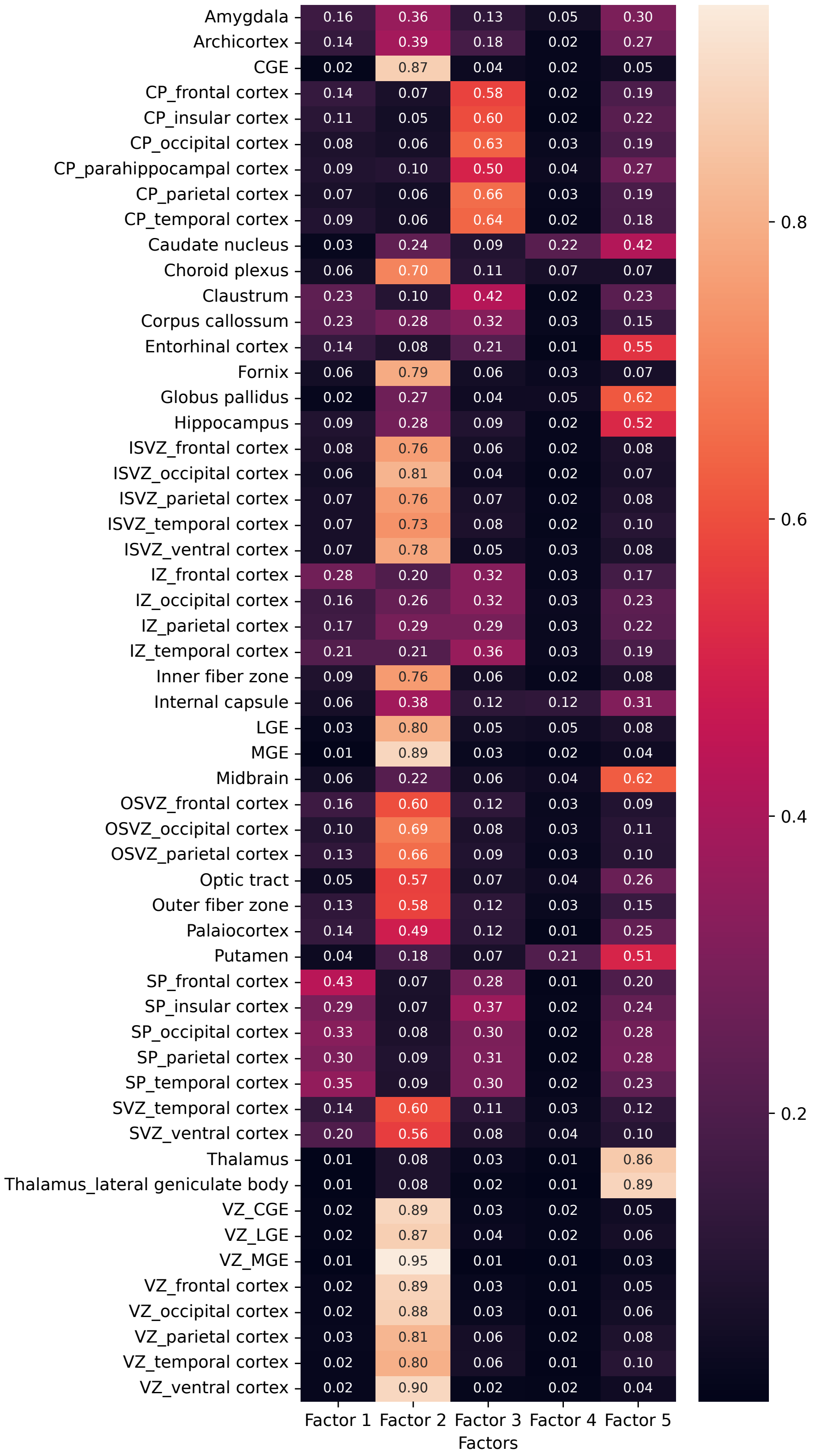


**Extended Data Fig. 9: Proportion of counts in each subregion explained by each factor.** The VZ of the medial ganglionic eminence (VZ_MGE) and the lateral geniculate nucleus (LGN) of the thalamus stand out in factor 2 and 5, having the largest proportion of their counts explained by each factor. Factor correspond to cortical subplate (factor 1), germinal zones in the cortex and the ganglionic eminences including ventricular and subventricular zones (factor 2), cortical plate (factor 3), caudate nucleus and putamen (factor 4) and thalamus (factor 5).

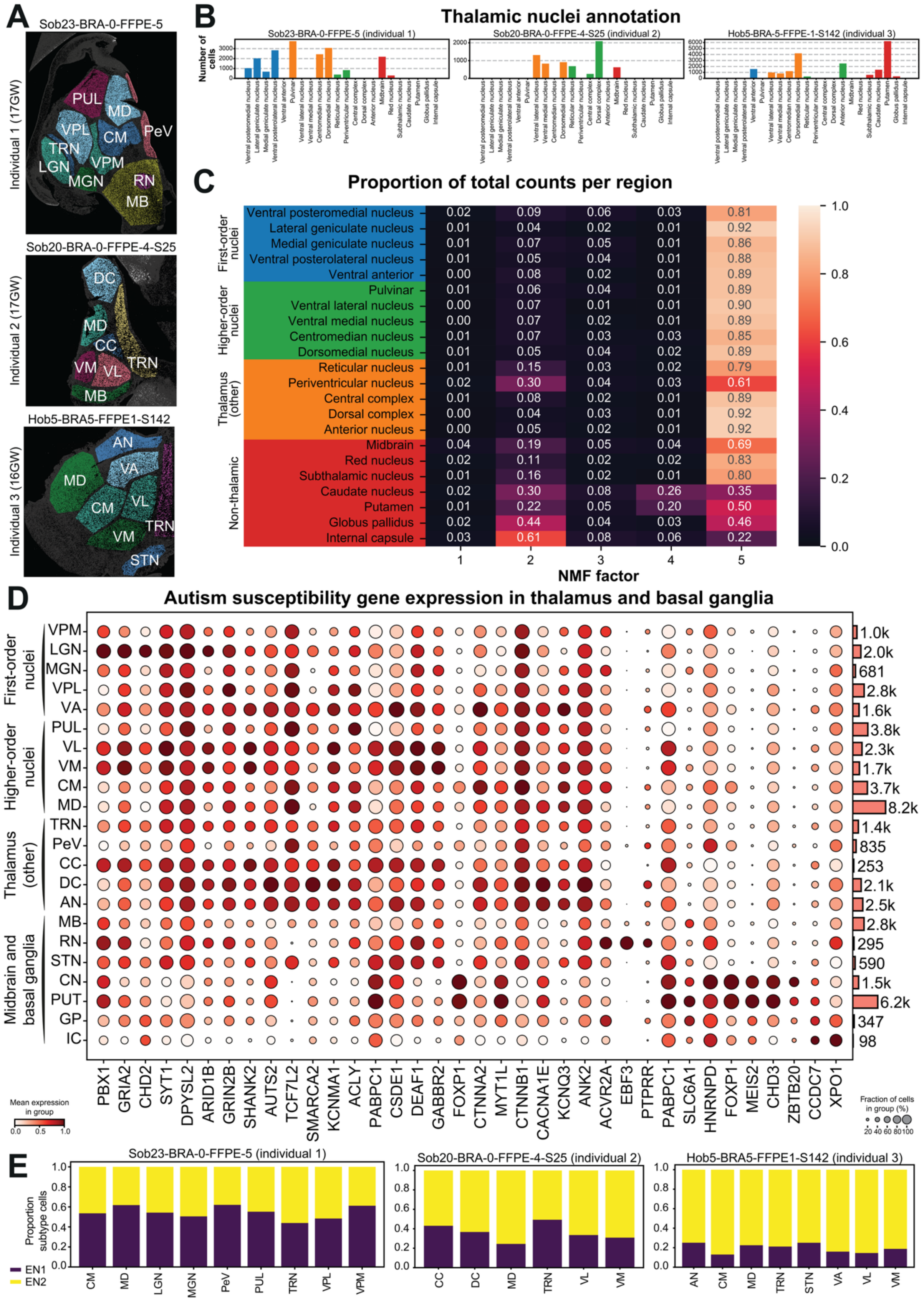


**Extended Data Fig. 10: Autism susceptibility gene expression in first-order and higher-order thalamic nuclei.**

***A)*** Spatial maps of annotated thalamic nuclei in three Xenium sections.

***B)*** Number of thalamic excitatory neurons (ENs) assigned to first-order (blue) or higher-order thalamic nuclei (orange), other thalamic regions (green) or midbrain and basal ganglia regions (red), shown per Xenium section.

***C)*** Proportion of total counts explained by each factor in thalamic excitatory neurons (mapped to reference thalamus single-cell data with Tangram) annotated to thalamic, midbrain and basal ganglia regions. Proportions were calculated using NMF cell loadings to annotated regions (**Methods**). Factor 5 (thalamus) explains most expression of thalamic regions, while factor 2 (germinal zones) and factor 4 (caudate nucleus and putamen) additionally explain a substantial proportion of non-thalamic gene expression.

***D)*** Dot plot showing mean log1p-normalised expression of selected autism susceptibility genes in annotated regions. VPM, Ventral posteromedial nucleus; LGN, lateral geniculate nucleus; MGN, medial geniculate nucleus; VPL, ventral posterolateral nucleus; VA, ventral anterior nucleus; PUL, pulvinar; VL, ventral lateral nucleus; VM, ventral medial nucleus; CM, centromedian nucleus; MD, dorsomedial nucleus; TRN, reticular nucleus; PeV, periventricular nucleus; CC, central complex; DC, dorsal complex; AN, anterior nucleus; MB, midbrain; RN, red nucleus; STN, subthalamic nucleus; CN, caudate nucleus; PUT, putamen; GP, globus pallidus; IC, internal capsule.

***E)*** Proportion of Tangram-classified EN subtypes in cells of annotated thalamic nuclei per Xenium section.

### Supplementary Note

**Enrichment of *de novo* mutations in regionally distinct autism predisposition genes.**

We investigated whether the rates of protein-truncating *de novo* mutations (DNMs) amongst robustly expressed, regionally distinct genes (programs) differ significantly between autistic children and children not diagnosed with autism. First, we collected DNMs observed in 250 autism predisposition genes ('Xenium panel genes') among 18,975 autistic children from several previously published cohorts[7,8,45]. We additionally evaluated DNMs in 7,881 siblings not diagnosed with autism from the same cohorts and 31,052 children primarily diagnosed with a developmental disorder (DD)[7,79]. The observed DNM rates in three regional programs (Factor 2: germinal zones; Factor 3: cortical plate; Factor 5: thalamus) are shown in **Fig. 2D (left)**. In all gene sets, the highest rate of DNMs was observed in probands ascertained for DDs, followed by those ascertained for autism, followed by their non-autistic siblings. Factors 1 (subplate, 3 genes) and 4 (caudate nucleus and putamen, 2 genes) were not tested separately due to their small size.

Next, we examined whether there is a significant difference in DNM enrichment between the main programs (factors) while accounting for mutation rates (**Fig. 2D, right**). We assigned each gene to the region with the highest loading; genes with highest relative loading on Factor 5 were considered thalamus-enriched genes (n=75), those with highest loading on Factor 2 were considered germinal-zones-enriched (n=36), whereas the remaining 8 genes from Factor 3 were enriched in the cortical plate. Two genes with equal loading on Factors 2 and 5 were assigned to the germinal zone genes.

We then tested the difference in enrichment between the three gene sets. Specifically, we used a Poisson test to examine whether the observed DNM rate ratio between any two groups (i.e. number DNMs in the first set divided by the number of DNMs in the second set) differs significantly from the expected ratio from a null model (i.e. the sum of gene-level mutation rates in the first gene set divided by the sum in the second set). We used Bonferroni correction to account for multiple testing (18 tests from 3 gene set comparisons of 2 DNM classes in 3 phenotypic groups; significant $p < 0.0028$).

**Fig. 2D (right)** shows the fold-difference between the observed over the expected rate ratios (i.e. relative risk of 1 means equal observed and expected rate ratios). Among autistic children, the observed protein-truncating DNM burden did not differ significantly between genes assigned to the cortical plates, the thalamus or the germinal zones ($p > 0.05$), whereas the burden of these DNMs in children ascertained for developmental disorders was significantly higher in the cortical plate (relative risk = 2.19; 95% CI = 1.72 to 2.76; $p = 1.0\times10^{-9}$) and germinal zone genes (relative risk = 1.52, 95% CI = 1.34 - 1.73, $p = 3.7\times10^{-11}$) *versus* thalamus genes. The observed rates of protein-truncating DNMs in siblings not diagnosed with autism as well as synonymous DNMs in all three groups (DD, ASD, siblings) were comparable across gene sets (**Supp. Note Fig. 1A**).

Last, we used recently published data on autism features in the Simons Simplex Collection (SSC) and Simons Foundation Powering Autism for Research Knowledge (SPARK) cohorts[80] to examine the association between protein-truncating DNMs in regionally distinct genes and factor scores capturing core autism features (F1: insistence on sameness; F2: social interaction; F3: sensory–motor behavior; F4: self-injurious behavior; F5: idiosyncratic repetitive speech and behavior; F6: communication skills). Here we used linear regression to assess the change in each factor score that is associated with having a protein-truncating DNM in thalamus-enriched genes vs. germinal zone genes, amongst 127 autistic children with *de novo* protein-truncating mutations in one of the robustly expressed genes (118 panel genes) and available factor scores. We used the scaled factor scores as an outcome, the affected gene set as a predictor and the gene mutation rate (log scaled) as well as the child's sex and age (inverse rank normalized) as covariates. There were no significant differences in factor scores associated with harboring a protein-truncating DNM in regionally distinct genes vs. other autism genes or in germinal zone genes vs. thalamus genes ($p > 0.05$) (**Supp. Note Fig. 1B**).

In summary, there was no evidence that de novo PTVs exert different degrees of predisposition to autism when their expression is enriched in the thalamus compared to the germinal zones (**Fig. 2D**). Note that this analysis estimates the average effect associated with a single DNM; given the larger number of genes that are robustly expressed in the thalamus vs. germinal zones, this implies that the cumulative risk at the population level is collectively higher in thalamus-enriched genes. Last, there was no evidence that core autism features differ depending on which of these two expression programmes the gene falls into. However, the sample size in this last analysis is very small (N=127) so it may be that it is simply underpowered.

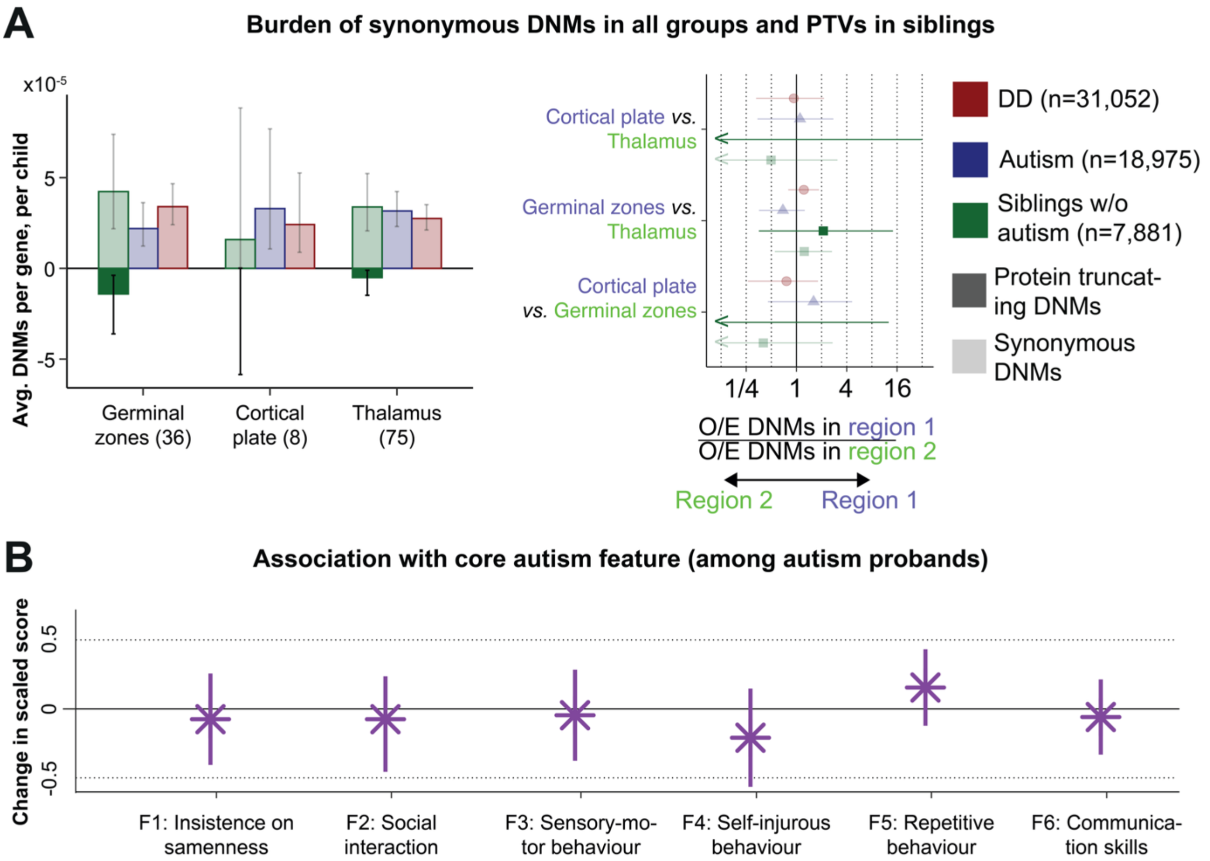


**Supp. Note Fig. 1: Enrichment of protein-truncating de novo mutations in regionally expressed autism predisposition genes and their association with core autism features.** ***A)*** Burden of synonymous de novo mutations (DNMs) in all groups and protein-truncating variants in siblings not diagnosed with autism. Overlapping genes were assigned to the region with higher factor loading, resulting in thalamus-enriched (n=75), germinal zones-enriched (n=36), and cortical plate-enriched (n=8) gene sets with different numbers than C. (Left) Average DNM burden per gene, per child in the gene sets across DD, autism and sibling cohorts. (Right) Pairwise differences in enrichment (observed DNM rate compared to expectation from underlying mutation rates) between brain regions. Arrows indicate the direction of enrichment to a brain region (coloured purple and green). P-values from two-sided Poisson test (** $p < 0.05$ after Bonferroni correction; * $p < 0.05$); error bars show 95%

confidence intervals. DD) developmental disorder cohort, O/E) observed over expected ratio. Synonymous DNMs are shown as a 'negative control'.

***B)*** Association with six core autism features (autism factor scores) amongst 127 autistic probands with de novo protein-truncating mutations in one of the robustly expressed genes (118 panel genes). Shown is the change in scaled score associated with having a protein-truncating DNM in thalamus-enriched genes vs. germinal zone genes as the outcome of a linear regression with the affected gene set as a predictor and the gene mutation rate (log scaled) as well as the child's sex and age (inverse rank normalized) as covariates. No significant differences in factor scores associated with harboring a protein-truncating DNM in regionally distinct genes vs. other autism genes or in germinal zone genes vs. thalamus genes were found ($p > 0.05$).
