## Supplementary Computational Note for "A spatial transcriptomic atlas of autism-associated genes identifies convergence in the developing human thalamus"

### Contents

**1 Xenium NMF model description 2**
1.1 Definition of subscripts and parameters 2
1.1.1 Subscripts 2
1.1.2 Observed model parameters 2
1.1.3 Free model parameters 2
1.2 Model Definition 2
1.3 Full Likelihood 3
1.4 Prior Distributions 3
1.5 Inference 5
1.5.1 Initial Value Computation 5
1.5.2 Model inference and choice of hyperparameters 5

1.6 Downstream Analysis . . . . . . . . . . . . . . . . . . . . . . . . . 6
1.6.1 Definition of Relative Cell and Gene Loadings . . . . . . . 6

### 1 Xenium NMF model description

#### 1.1 Definition of subscripts and parameters

##### 1.1.1 Subscripts

- cell index $c \in \{1 \dots C\}$
- gene index $g \in \{1 \dots G\}$
- region index $r \in \{1 \dots R\}$
- factor index $f \in \{1 \dots F\}$

##### 1.1.2 Observed model parameters

- Observed counts in each cell $X_{cg}$
- One hot assignment of cells to anatomical brain regions $H_{cr}$
- One hot assignment of cells to experimental batch (corresponding to sections in Xenium experiments) $H_{ce}$

##### 1.1.3 Free model parameters

- expected counts $x_{cg}$
- cell loadings of each factor $w_{cf}$
- gene loadings of each factor $h_{fg}$
- detection efficiency of genes in each cell $l_c$
- ambient RNA (e.g. "background" binding) of genes in each experimental batch $s_{eg}$

#### 1.2 Model Definition

Non-negative matrix factorization decomposes the matrix of biological expression value $x_{cg}^{(B)}$ into two smaller matrices of dimensions cells x factors and factors x genes [1, 2]:

$$x_{cg}^{(B)} = \sum_{F} w_{cf} h_{fg} \tag{1}$$

We take into account technical factors of variation that change the expectation values for measured quantitites of counts by transforming "biological" expectation values to account for technical variables:

$$x_{cg}^{(M)} = l_c * (H_{ce} * s_{eg} + x_{cg}^{(B)}) \tag{2}$$

Here, $H_{ce}$ denotes a one-hot categorical assignment of cells to experimental batches - which correspond to individual sections in Xenium data - and $l_c$ describes differences in detection efficiency of genes across cells and $s_{ge}$ models ambient RNA (e.g. "background binding") for each gene in each batch. These "measurement" expectation values are then used to parameterize a Negative Binomial observation model of the observed raw count values $X_{cg}$:

$$X_{cg} \sim \texttt{NegBinom}(\mu = x_{cg}^{M}, \alpha = a_g) \tag{3}$$

Here, $a_g$ are Negative Binomial over-dispersion parameters for each gene.

#### 1.3 Full Likelihood

In summary, the full likelihood of the model is:

$$\prod_C \prod_G \prod_J P(X_{cg}|\theta_{cg}) = \prod_C \prod_G \prod_J \texttt{f}_{\texttt{NB}}(\mu = F(\theta_{cg}), \alpha = a_g), \tag{4}$$

where $\theta_{cg}$ are all model parameters:

$$\theta_{cg} = (l_{cg}, H_{ce}, s_{eg}, h_{fg}, w_{cf}) \tag{5}$$

, $\texttt{f}_{\texttt{NB}}$ is the probability density function of the Negative Binomial distribution and $F$ is a function that summarizes the dependence of expected measured counts $x_{cg}^{M}$ on all model parameters, through equations 1, 2.

#### 1.4 Prior Distributions

The full Bayesian posterior of the model, $P(\theta_{cg}|X_{cg})$, is obtained by combining the likelihood above, $P(X_{cg}|\theta_{cg})$, with the prior distributions for each parameter, $P(\theta_{cg})$:

$$P(\theta_{cg}|X_{cg}) = P(X_{cg}|\theta_{cg})P(\theta_{cg}) \tag{6}$$

We list and motivate these prior distributions below.

We use a hierarchical prior for $w_{cf}$:

$$w_c^{(0)} \sim \texttt{Gamma}(\alpha = 2.0, \beta = 2.0) \tag{7}$$

$$w_{cf} \sim \texttt{Gamma}(\alpha = \frac{w_c^{(0)}}{F}, \beta = F) \tag{8}$$

$w_c^{(0)}$ can be interpreted as the number of factors that are active in each cell.

Priors for $h_{hg}$ are given by:

$$h_g \sim \texttt{Gamma}(\alpha = 1, \beta = 1) \tag{9}$$

$$h = 3 \tag{10}$$

$$h_{fg} \sim \texttt{Gamma}(\alpha = \frac{h}{F}, \beta = \frac{1}{h_g}) \tag{11}$$

$h_g$ can be interpreted as the total counts produced by a gene in a cell per factor in the steady state. $h$ can be interpreted as the number of factors this gene is part of, since the mean per factor is given by:

$$\texttt{mean}(h_{fg}) = \frac{h \cdot h_g}{F} \tag{12}$$

Moving on to technical variables, we use this hierarchical prior for the overdispersion parameter:

$$a_g = \frac{1}{E_g^2} \tag{13}$$

$$E_g \sim \texttt{Exponential}(\phi) \tag{14}$$

$$\phi \sim \texttt{Gamma}(\mu = 3, \sigma = 1) \tag{15}$$

This kind of prior is called a containment prior [3]. As also shown in [4], to understand the reasoning behind it, it is useful to write the total variance of the Negative Binomial distribution:

$$\sigma^2 = \mu + \frac{\mu^2}{a}, \tag{16}$$

so the chosen prior pushes alpha towards infinity and with that the "extra variance" compared to a Poisson distribution towards 0. So inference should only converge to solutions with considerable overdispersion if this is really necessary.

The mean relative detection probability in each cell depends on the experimental batch $e$ each cell comes from:

$$l_e \sim \texttt{Beta}(\alpha = 1, \beta = 1) \tag{17}$$

(This results in a broad distribution between 0 and 1 with mean of 0.5.)

$$l_c \sim \texttt{Gamma}(\alpha = 10, \beta = \frac{10}{H_{ce} * l_e}) \tag{18}$$

Finally, we use this containment prior for the ambient RNA parameter:

$$s_e \sim \texttt{Gamma}(\mu = 0.005, \sigma = 0.00005) \tag{19}$$

$$b_e \sim \texttt{Exponential}(\lambda^b = 9) \tag{20}$$

$$a_e^s = \frac{1}{b_e^2} \tag{21}$$

$$s_{eg} \sim \texttt{Gamma}(\alpha = a_e^s, \beta = \frac{a_{ej}^s}{s_e}) \tag{22}$$

#### 1.5 Inference

##### 1.5.1 Initial Value Computation

All parameter initial values, except the cell loadings, are set to the mean of their prior distributions. For cell loadings, we use a fast PCA-based heuristic implemented in the `find_initial_values` function of our `Xenium_NMF` python package. This is similar to other widely used factorization packages (e.g. MOFA+ [5]) that use PCA components as initial values for factors to improve the speed and reproducibility of inference. Our heuristic for setting initial values for the cell loadings has three steps:

1. **Dimensionality Reduction:** Principal component analysis (PCA) is applied to the data.
2. **Gene Clustering:** Genes are clustered based on their PC loadings.
3. **Initial Loading Calculation:** Initial cell loadings are calculated as the average gene expression values of gene clusters.

##### 1.5.2 Model inference and choice of hyperparameters

The model is implemented in the pebrobabilistic programming language pyro [6]. Inference is performed using stochastic variational inference (SVI), using the "AutoHierarchical NormalMessenger" autoguide and Adam optimizer [7] for 40000 iterations with a learning rate of 0.001 and a batch size of 2171772 cells. This batch size simply corresponds to the maximal number of cells that fit into GPU memory on a Tesla V100-SXM2 32GB GPU. The median of the posterior distribution is then extrated from the variational distribution to give point estimates for $w_{cf}$ and $h_{fg}$.

In more detail, within the SVI inference framework, the posterior distributions over unknown parameters are approximated by univariate normal distributions that are transformed to ensure the appropriate scale for each parameter (for example, positive for the mean of a Gamma distribution). The parameters of these variational distributions are then determined by minimizing the KL divergence between the variational approximation and the true posterior distribution or, equivalently, maximizing the evidence lower bound (ELBO loss function). The use of the "AutoHierarchical NormalMessenger" autoguide improves the variational approximation, by keeping dependencies between parameters as present in the model through hierarchical prior distributions. Specifically, the mean-field posterior at any site is a transformed normal distribution, the mean of which depends on the value of that site given its dependencies in the model:

$$\underbrace{x}_{\text{Variable}} = \texttt{transform}(\texttt{Normal}(\underbrace{\mu}_{\text{Total mean}}, \underbrace{\sigma}_{\text{Independent standard deviation}})), \tag{23}$$

$$\underbrace{\mu}_{\text{Total mean}} = \underbrace{loc}_{\text{Independent mean}} + \underbrace{\texttt{transform.inverse}(\hat{\mu})}_{\text{Prior mean transformed to real value space}} \cdot \underbrace{weight}_{\text{Weight}}, \tag{24}$$

which is equivalent to defining a posterior distribution conditional on the hierarchical prior of each variable, where `transform` indicates a function that maps the posterior distribution in real value space to the domain in which the variable is defined.

#### 1.6 Downstream Analysis

##### 1.6.1 Definition of Relative Cell and Gene Loadings

Since the expected gene expression is given by:

$$\hat{x}_{cg} = \sum_{f=1}^{F} w_{cf} h_{fg}, \tag{25}$$

the expected expression from a single factor $f'$ is

$$\hat{x}_{cf'g} = w_{cf'} h_{f'g} \quad \text{for } f' \in \{1, \dots, F\}. \tag{26}$$

The *relative cell loading* of a factor $f'$ is defined as the proportion of total counts in a cell that this factor explains:

$$\hat{x}_{cf'} = \frac{\sum_g \hat{x}_{cf'g}}{\sum_g \hat{x}_{cg}}. \tag{27}$$

The *total cell loading* for factor $f'$ in each cell is defined as the total number of counts that the factor explains:

$$\hat{x}_{cf'}^{(\text{Total})} = \sum_g \hat{x}_{cf'g}. \tag{28}$$

The *average relative cell loading* in a region is then given by:

$$\hat{x}_{rf'} = \frac{\sum_c \left( H_{cr} \frac{\sum_g \hat{x}_{cf'g}}{\sum_g \hat{x}_{cg}} \right)}{\sum_c H_{cr}}, \tag{29}$$

and the *average total cell loading* in a region is given by:

$$\hat{x}_{rf'}^{(\text{Total})} = \frac{\sum_c \left( H_{cr} \sum_g \hat{x}_{cf'g} \right)}{\sum_c H_{cr}}, \tag{30}$$

where $H_{cr}$ represents the weight (or membership indicator) of cell $c$ in region $r$.

We combine existing regions into a smaller set of factor-associated regions $P_{f'}$ by including all regions in $R$ that have an average total cell loading greater than the expectation value across regions, i.e.,

$$P_{f'} = \bigcup_{r \in I_{f'}} R_r \quad \text{where} \quad I_{f'} = \left\{ r \mid \hat{x}_{rf'}^{(\text{Total})} > E\left(\hat{x}_{rf'}^{(\text{Total})}\right) \right\}, \tag{31}$$

and

$$E\left(\hat{x}_{rf'}^{(\text{Total})}\right) = \frac{1}{|R|} \sum_{r \in R} \hat{x}_{rf'}^{(\text{Total})}. \tag{32}$$

The proportion of a gene's counts in a cell $c$ that is explained by factor $f'$ is given by:

$$\hat{x}_{f'gc} = \frac{\hat{x}_{cf'g}}{\hat{x}_{cg}}, \tag{33}$$

and the *relative gene loading* is defined as the average proportion of counts in each cell that is explained by factor $f'$ across all regions in $P_{f'}$ for a particular gene:

$$\hat{x}_{f'g} = \frac{1}{|P_{f'}|} \sum_{p \in P_{f'}} \frac{\sum_c \left( H_{cp} \frac{\hat{x}_{cf'g}}{\hat{x}_{cg}} \right)}{\sum_c H_{cp}}, \tag{34}$$

where $H_{cp}$ denotes the weight (or membership indicator) of cell $c$ in region $p$.

Values of $\hat{x}_{f'g}$ larger than $\frac{1}{F}$ show higher than average loading for a gene in a factor. To extract only *factor-enriched* genes, we set a threshold $s$ that goes beyond this average:

$$s = \frac{1}{F-2} \tag{35}$$

, where $F$ is the number of factors. For 5 factors this results in a threshold of 0.33 to denote factor enriched genes.
